# Identification and prediction of Parkinson’s disease subtypes and progression using machine learning in two cohorts

**DOI:** 10.1101/2022.08.04.502846

**Authors:** Anant Dadu, Vipul K. Satone, Rachneet Kaur, Sayed Hadi Hashemi, Hampton Leonard, Hirotaka Iwaki, Mary B. Makarious, Kimberley Billingsley, Sara Bandres-Ciga, Lana J. Sargent, Alastair J. Noyce, Ali Daneshmand, Cornelis Blauwendraat, Ken Marek, Sonja W. Scholz, Andrew B. Singleton, Mike A. Nalls, Roy H. Campbell, Faraz Faghri

## Abstract

**Background:** The clinical manifestations of Parkinson’s disease (PD) are characterized by heterogeneity in age at onset, disease duration, rate of progression, and the constellation of motor versus non-motor features. There is an unmet need for the characterization of distinct disease subtypes as well as improved, individualized predictions of the disease course. The emergence of machine learning to detect hidden patterns in complex, multi-dimensional datasets provides unparalleled opportunities to address this critical need.

**Methods and Findings:** We used unsupervised and supervised machine learning methods on comprehensive, longitudinal clinical data from the Parkinson’s Disease Progression Marker Initiative (PPMI) (n = 294 cases) to identify patient subtypes and to predict disease progression. The resulting models were validated in an independent, clinically well-characterized cohort from the Parkinson’s Disease Biomarker Program (PDBP) (n = 263 cases). Our analysis distinguished three distinct disease subtypes with highly predictable progression rates, corresponding to slow, moderate, and fast disease progression. We achieved highly accurate projections of disease progression five years after initial diagnosis with an average area under the curve (AUC) of 0.92 (95% CI: 0.95 ± 0.01 for the slower progressing group (PDvec1), 0.87 ± 0.03 for moderate progressors, and 0.95 ± 0.02 for the fast progressing group (PDvec3). We identified serum neurofilament light (Nfl) as a significant indicator of fast disease progression among other key biomarkers of interest. We replicated these findings in an independent validation cohort, released the analytical code, and developed models in an open science manner.

**Conclusions:** Our data-driven study provides insights to deconstruct PD heterogeneity. This approach could have immediate implications for clinical trials by improving the detection of significant clinical outcomes that might have been masked by cohort heterogeneity. We anticipate that machine learning models will improve patient counseling, clinical trial design, allocation of healthcare resources, and ultimately individualized patient care.

## Introduction

Parkinson’s disease (PD) is a complex, age-related neurodegenerative disease that is defined by a combination of core diagnostic features, including bradykinesia, rigidity, tremor, and postural instability (Hughes et al. 1992; Postuma et al. 2015). Substantial phenotypic heterogeneity is well recognized within the disease, complicating the design and interpretation of clinical trials and limiting patients’ counseling about their prognosis. The clinical manifestations of PD vary by age at onset, rate of progression, associated treatment complications, as well as the occurrence and constellation of motor/nonmotor features.

The phenotypic heterogeneity that exists within the PD population poses a major challenge for clinical care and clinical trial design. A clinical trial has to be suitably powered to account for interindividual variability, and as a consequence, trials are either large, long, expensive, and/or only powered to see large effects. This problem becomes particularly burdensome as we move increasingly towards early-stage trials when therapeutic interventions are likely to be most effective. To that effect, defining subcategories of PD and the ability to predict even a proportion of the disease course has the potential to significantly improve cohort selection, inform clinical trial design, reduce the cost of clinical trials, and increase the ability of such trials to detect treatment effects.

Attempts thus far at the characterization of disease subtypes have followed a path of clinical observation based on age at onset or categorization based on the most observable features (Stebbins et al. 2013). Thus, the disease is often separated into early-onset versus late-onset disease, slowly-progressing “benign” versus fast-progressing “malignant” subtypes, PD with or without dementia, or based on the most prominent clinical signs into a tremor-dominant versus a postural instability with gait disorder subtype (Jankovic et al. 1990; Zetusky, Jankovic, and Pirozzolo 1985). This dichotomous separation, while intuitive, does not faithfully represent the clinical features of the disease, which are quantitative, complex, and interrelated. A more realistic representation of the disease and disease course requires a transition to a data-driven, multi-dimensional schema that encapsulates the constellation of interrelated features and allows tracking (and ultimately predicting) change.

Previous studies used cluster analysis, a data-driven approach, to define two to three clinical PD subtypes (van Rooden et al. 2010; Fereshtehnejad et al. 2015, 2017; Lawton et al. 2018). Depth of phenotypic information and longitudinal assessments in these studies were variable and often limited to certain clinical features and short-term follow-ups. Moreover, many previous studies were limited by insufficient methods to capture longitudinal changes over multiple assessment visits. To this date, none of the previous approaches to PD clustering were replicated in an independent cohort with transparent code and analysis.

We have previously used multi-modal data to produce a highly accurate disease status classification and to distinguish PD-mimic syndromes from PD (Nalls et al. 2015). These efforts demonstrated the utility of data-driven approaches in the dissection of complex traits and have also led us to the next logical step in disease prediction: supplementing the prediction of whether a person has or will have PD also to include a prediction of the timing and directionality of the course of their disease.

Here, we describe our work on delineating and predicting the clinical progression of PD. The first stage of this effort requires creating a multi-dimensional space that captures the disease’s features and the progression rate of these features (i.e., velocity). Rather than creating a space based on *a priori* concepts of differential symptoms, we used data dimensionality reduction methods on the complex clinical features observed 60 months after initial diagnosis to create a meaningful spatial representation of each patient’s status at this time point. After creating this space, we used unsupervised clustering to determine whether there were clear subtypes of disease within this space. This effort identified three distinct clinical subtypes corresponding to three groups of patients progressing at varying velocities (i.e., slow, moderate, and fast progressors). These subtypes were validated and replicated in an independent cohort. Following the successful creation of disease subtypes within a progression space, we created a baseline predictor that accurately predicted an individual patient’s clinical group membership five years later. Further, we examined the predictive capability of biospecimen biomarkers at baseline and the genetic information in identifying the subtypes. Our work highlights the utility of machine learning as an ancillary diagnostic tool to identify disease subtypes and project individualized progression rates based on model predictions.

## Methods

### Study design and participants

This study included clinical data from the Parkinson’s Progression Marker Initiative (PPMI, http://www.ppmi-info.org/) and the Parkinson’s Disease Biomarkers Program (PDBP, https://pdbp.ninds.nih.gov/). Both cohort’s data went through triage for missing data, 60-month assessment (36-month in PDBP), and comprehensive phenotype collection. Only data from participants with 60 months of follow-up for PPMI and 36 months for PDBP were included in the study. Overall, in the PPMI (n = 294 PD cases including 99 (34%) female; 154 controls including 58 (38%) female), and in the PDBP (n = 263 PD cases including 112 (43%) female; 115 controls including 64 (56%) female) passed the triage. The PPMI average age at the screening of PD cases was 61 ± 9.7 years and 60.3 ± 11 years for controls. The PDBP average age of PD cases was 64.3 ± 8.6 years and 63.6 ± 9.5 years for controls. The PPMI data also included 28 patients with other enrollments (10 PRODROMA; 8 GENPD; 6 GENUN; 3 SWEDD; 1 REGPD), which were excluded. The PPMI and PDBP cohorts consist of observational data from comprehensively characterized PD patients and matched controls. All PD patients fulfilled the UK Brain Bank Criteria (Sudlow et al. 2015). Control subjects had no clinical signs suggestive of parkinsonism, no evidence of cognitive impairment, and no first-degree relative diagnosed with PD. Age and MDS-UPDRS Part III (objective motor symptom examination by a trained neurologist) distribution of cohorts at baseline were investigated using Kernel Density Estimation (KDE) to show that these independent cohorts are identically distributed and ensure the integrity of replication and validation. Each contributing study abided by the institutional review boards’ ethics guidelines. All participants gave informed consent for inclusion in their initial cohorts and subsequent studies. **Figure 1** provides an overview of the analyses and study design.

**Fig 1.**
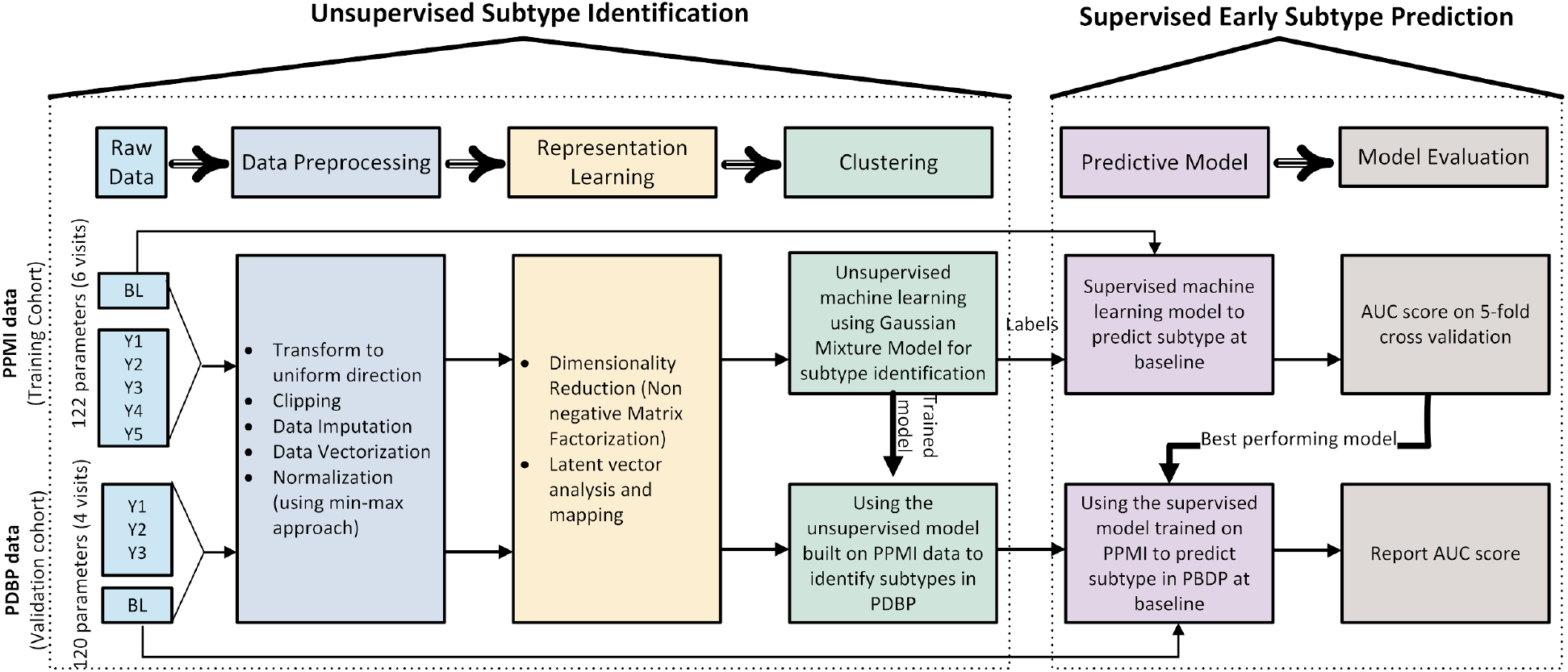
Workflow of analysis and model development.

### Dataset Construction

The discovery and replication cohorts include visit data collected every 12 months starting from baseline to 60 months (36 months for PDBP) follow-up. In PPMI, visits at the 6 and 9-month time points from baseline were excluded in our analysis due to the high data missingness rate (>50%).

For each cohort, a comprehensive and shared set of longitudinally collected common clinical data elements were selected for analysis. Overall, 122 clinical features were available across six visits for PPMI (Table S1) and 120 features across four visits for PDBP. We used the following features for the subtype identification stage:

i. International Parkinson’s disease and Movement Disorder Society Unified Parkinson’s Disease Rating Scale (MDS-UPDRS) Part I, Part II, and Part III (Goetz et al. 2008)
ii. Cranial Nerve Examination (CN I-XII)
iii. Montreal Cognitive Assessment (Nasreddine et al. 2005)
iv. Hopkins Verbal Learning Test (Brandt 1991)
v. Semantic Fluency test (Goodglass, Kaplan, and Barresi 2001)
vi. WAIS-III Letter-Number Sequencing Test (Wechsler 1997)
vii. Judgment of Line Orientation Test (Benton, Varney, and Hamsher 1978)
viii. Symbol Digit Modalities Test (SMITH and A 1968)
ix. SCOPA-AUT (Visser et al. 2004)
x. State-Trait Anxiety Inventory for Adults (Spielberger et al. 1983)
xi. Geriatric Depression Scale (Yesavage and Sheikh 1986)
xii. Questionnaire for Impulsive-Compulsive Disorders in Parkinson’s Disease (Weintraub et al. 2009)
xiii. REM-Sleep Behavior Disorder Screening Questionnaire (Stiasny-Kolster et al. 2007)
xiv. Epworth Sleepiness Scale (Johns 1991).

In addition to these clinical measurements, biological and genetic-based features were included in the baseline subtype interpretation and the subtype prediction. These additional features include genotypes using imputed Illumina NeuroX array, vital signs, serum, CSF, and urine measurements. For genotyping data, we used the variants mapped to human genome build 38 (hg38) genotyping from unrelated European ancestry imputed genotype data passing standard QC metrics used to construct the genetic risk score (GRS) in Nalls et al 2019. The values indicate the number of copies of the minor allele of each variant for each subject. We used these values as categorical features. Patient characteristics include height, weight, blood pressure, and demographic details. For biological biomarkers, we assessed alpha-synuclein, total tau protein, β-amyloid 1–42 (Aβ42), phospho-tau181 (p-Tau181) in CSF, serum neurofilament light (NfL), and urine levels of di-22:6-bis (monoacylglycerol) phosphate total in the urine. We studied the longitudinal variation of biomarkers and patients’ characteristics measurements across the identified PD subtypes. Furthermore, we investigated the biological measurements’ role in discriminating PD subtypes. Please refer to the **Online Methods** in the **Data Preprocessing** section for details on data preprocessing.

### Procedures and statistical analysis

The data analysis pipeline for this work was performed in Python (version 3.8) with the support of several open-source libraries (NumPy, pandas, matplotlib, seaborn, plotly, scikit-learn, UMAP, XGBoost, LightGBM, H2O, streamlit). To facilitate replication and expansion of our work, we have made the notebook publicly available on GitHub at https://github.com/anant-dadu/PDProgressionSubtypes. The code is part of the supplemental information; it includes the rendered Jupyter notebook with full step-by-step data preprocessing, statistical, and machine learning analysis. For readability, machine learning parameters have been described in the Python Jupyter notebook and not in the text of the paper. Our results are available on an interactive web browser (https://anant-dadu-pdprogressionsubtypes-streamlit-app-aaah95.streamlitapp.com/), which allows users to browse the PD progression space. In addition, the browser also includes predictive model interpretations allowing readers to explore feature contributions to model performance.

### Unsupervised subtype identification

We used dimensionality reduction techniques to develop an interpretable representation of high modality longitudinal data. Dimensionality reduction techniques helped us to build the “progression space” where we can approximate each patient’s position relative to both controls and other cases after the 60-month period in one-year intervals. We used the Non-negative Matrix Factorization (NMF) technique to achieve this aim (D. D. Lee and Seung 1999; Daniel D. Lee and Seung 2001). Alternative methods, such as principal component analysis and independent component analysis, did not perform as well as NMF on longitudinal clinical data due to the non-negative nature of our clinical tests. This process essentially collapses mathematically related parameters into the same multi-dimensional space, mapping similar data points close together. Details on the NMF procedure and its latent space adjustment can be found in the **Online Methods** - **NMF** and **Latent Space Adjustment** sections.

### Supervised early subtype prediction

After identifying progression classes using unsupervised learning, we built predictive models utilizing multiple supervised machine learning methods, including the *ensemble learning* approach. This method combines multiple learning algorithms to generate a better predictive model than could be obtained using a single learning algorithm (Rokach 2010). To do this, we used stacking ensembles of three supervised machine learning algorithms (Random forest (Breiman 2001), LightGBM (Ke et al. 2017), and XGBoost (Chen and Guestrin 2016)) to predict PD clinical subtypes using the data obtained at the time a neurologist first reviewed the patient as the input (combining baseline and varied time points). This approach outperformed other methods in preliminary testing, such as support vector machines (SVM) and simple lasso-regression models. Besides the predictive performance, we chose an ensemble approach due to the nature of our data and problem: (i) decision trees are intrinsically suited for multiclass problems, while SVM is intrinsically two-class, (ii) they work well with a mixture of numerical, categorical, and various scale features, (iii) they can be used to rank the importance of variables in a classification problem and in a natural way which helps the interpretation of clinical results, and (iv) it also gives us the probability of belonging to a class, which is very helpful when dealing with individual subject progression prediction.

We developed three predictive models to predict the patient’s progression class after 60 months based on varying input factors: (a) from baseline clinical factors, (b) from baseline and first-year clinical factors, and (c) from biomarkers and genetic measurements. Please refer to the **Online Methods** in the **Model Evaluation and Tuning** section for additional information and a summary of model evaluation and tuning.

To validate the effectiveness of our predictive models, we randomly split the dataset into training (80%) and test (20%) sets. The training set was used for hyperparameter tuning and model training, while the test set is used to evaluate the model performance. We followed a 5-fold cross-validation (CV) approach for the hyperparameter tuning of model parameters. Specifically, we randomly divided the training set into five subsamples (folds). Each of the subsamples used exactly once as the validation data. The hyperparameters were chosen based on their average performance during cross-validation. The final performance is reported on the holdout test set. We repeated the whole procedure five times to get better estimates of model performance. The workflow of the approach is depicted in **Figure S1**.

The performance of the algorithm was measured by the area under the receiver operating curve (AUC) generated by plotting sensitivity vs. (1 − specificity). We used a macro-average AUC score computed by averaging the metric independently for each class (hence treating all classes equally for predicting fast, moderate, and slow progressing cases). The five results from the multiple iterations were averaged to produce a single estimation of performance across these three classes.

To conclusively validate the algorithm, we also evaluated the performance of the predictive models (trained on the PPMI measurements) on the independent PDBP cohort. To replicate, we trained the supervised model on PPMI latent weights (at baseline) and then used the same model on the PDBP latent weights (at baseline). We show that the predictive models preserve their high accuracy applied to another dataset.

## Results

### Clustering vectors of progression

Figure 2 shows the result of the mathematical projection of PD progression, called *Parkinson’s disease progression space* detailing normalized progression trajectories of each sample relative to others based on this unsupervised classification system. This space shows the relative progression velocity of each patient in 60 months (i.e., speed and direction). The progression velocity level is divided into three main dimensions: motor, cognitive, and sleep-related disturbances. Based on latent variables clustered within the Parkinson’s progression space, the projected motor dimension was responsible for 63.58% of the explained variance, followed by the sleep dimension (21.81%), and cognitive dimension (14.61%). Across these trajectories, the unsupervised learning analysis reveals and classifies patients into three main subtypes of PD, relating to rates of disease progression: slow progressors (PDvec1), moderate progressors (PDvec2), and fast progressors (PDvec3). This shows how we can map the clinical features and progression velocity from the point of diagnosis. The components of the motor, cognitive and sleep dimensions can be found in the **Online Results** with a discussion of the latent space used to define the progression space that may aid in interpretability, as well as a summary in **Figure S2**.

**Fig 2.**
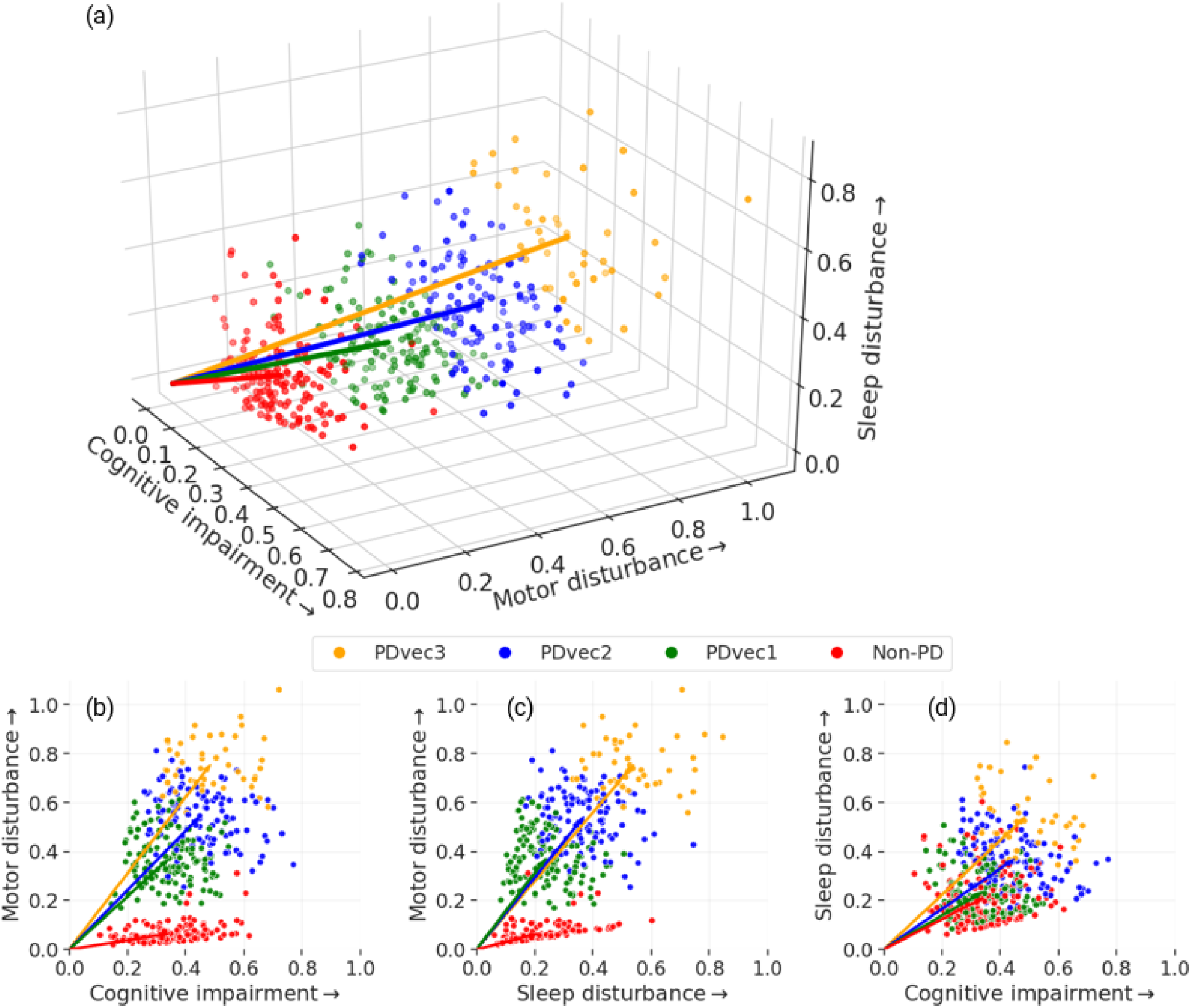
Different views of the Parkinson’s disease progression space in five years with three corresponding projected dimensions (cognitive, motor, and sleep dimensions) on a normalized scale. Subtypes of PD are identified using unsupervised learning (PDvec1, PDvec2, and PDvec3). (a) shows the view of all three dimensions, (b) view of the motor and cognitive dimensions, (c) view of motor and sleep dimensions, and (d) view of sleep and motor dimensions.

### Identified Subtypes and their Clinical Characteristics

Figure 3 shows the visualization of unsupervised learning via GMM in a two-dimension progression space. In two-dimensional progression space, the projected dimensions represent motor (y-axis) and cognitive (combined with sleep) (x-axis) components. Projected dimensions are normalized; the increase in values along either direction signifies a higher decline. GMM fits the data into different subtypes relating to velocity of decline across symptomatologies from non-PD controls. The Bayesian information criterion has identified three Gaussian distributions representing three PD subtypes. These three groups identified algorithmically within the case population change over time differently within the progression space and across specific biomarkers of progression, with PDvec3 generally progressing at a much steeper slope. Details of this can be seen in the **Online Results** section describing the **Five-Year PD Progression Space**.

**Fig 3.**
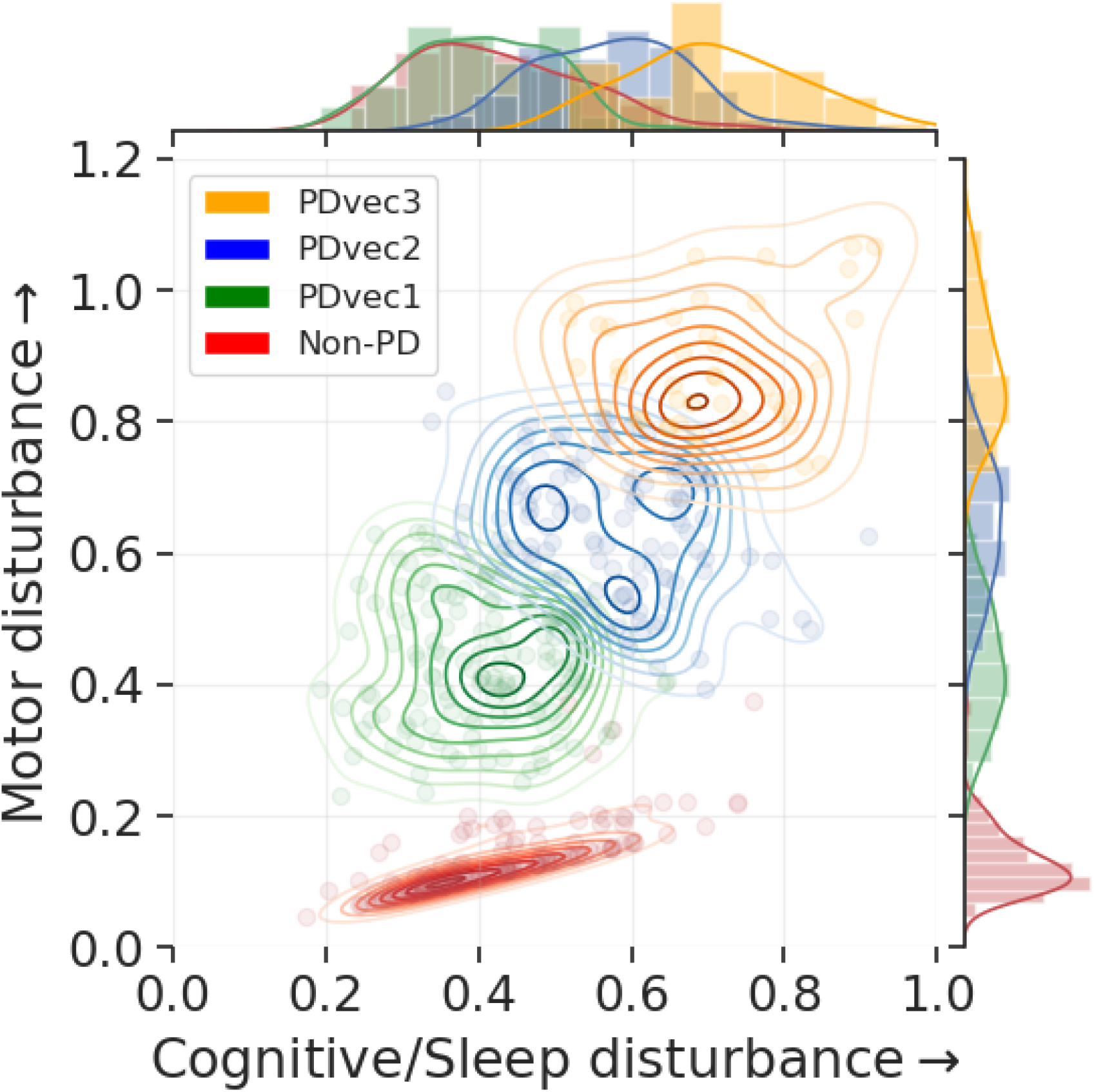
PD five-year progression space. Visualization of unsupervised learning via GMM on two-dimensional progression space and identification of three Gaussian distributions representing three distinct PD subtypes. An increase in value along either direction reflects the increase in the disturbance on a normalized scale.

### Biological Characteristics of the Identified Subtypes

Figure 4 shows the variation of biological biomarkers for each PD subtype over time. In terms of patients’ features, height and weight show a significant decline over time for the fast progressors (PDvec3) compared to other subtypes. In terms of blood biomarkers, NfL values are significantly higher after five years for the fast progressors in PDvec3 compared to PDvec1, with a steeper slope across time as well as higher values at baseline (P < 0.005 for both). Detailed results of Nfl comparisons across PDvec1, PDvec2 and PDvec3 can be found in **Table S4**. PD patients have lower values compared to healthy controls for all CSF sample measurements such as alpha-synuclein, total tau protein, Aβ42, and p-Tau181.

**Fig 4.**
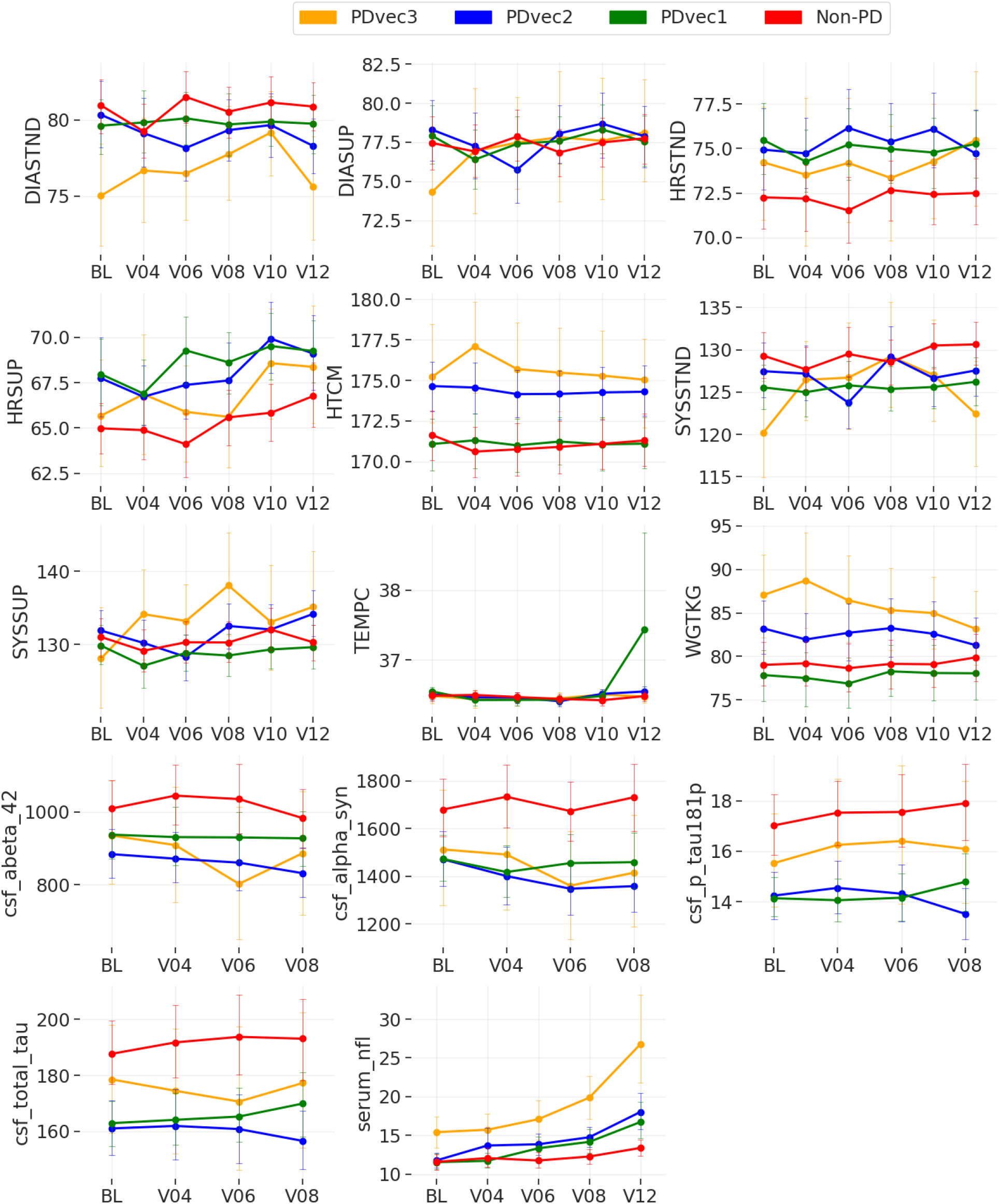
Shows the biological biomarker variation of each PD subtype over time. The graphs demonstrate the actual clinical values of each subtype overtime for vital signs (DIASTND: standing diastolic blood pressure (BP), DIASUP: supine diastolic BP, HRSTND: standing heart rate, HRSUP: supine heart rate, SYSSTND: standing systolic BP, SYSSUP: supine systolic BP, HTCM: height in cm, TEMPC: temperature in C, WGTKG: weight in kg), cerebrospinal fluid (abeta_42: β-amyloid 1–42, alpha_syn: alpha-synuclein, p_tau181p: phospho-tau181, total_tau: total tau protein), and serum neurofilament light levels (serum_nfl). BL: Baseline. V04: visit number 4 after 12 months. V06: visit number 6 after 24 months. V08: visit number 8 after 36 months. V10: visit number 10 after 48 months. V12: visit number 12 after 60 months.

### Genetic Analysis of the Identified Subtypes

In terms of the genetic association of PDs identified subtypes, genetic risk scores (GRS) were calculated as per (Nalls et al. 2019). As a one-time measurement, the GRS was not included during the longitudinal clustering exercise; however, we analyzed regressions comparing associations between the GRS and either the continuous predicted cluster membership probability (linear regression) or the binary membership in a particular cluster group compared to the others. All models were adjusted for age at onset, biological sex, and principal components as covariates to adjust for population substructure in PPMI. The GRS was significantly associated with decreasing magnitude of the sleep vector per Standard deviation (SD) of increase in the GRS (beta = -0.029, se = 0.010, p = 0.002, adjusted r2 = 0.046). For binary models of membership, we see that the GRS is weakly but significantly associated with a decreased risk of membership in PDvec3 (odds ratio = 0.563 per 1 SD increase from case GRS mean, beta = -0.574, se = 0.244, P = 0.018) and increased risk of membership in PDvec1 (odds ratio = 1.341, beta = 0.293, se = 0.134, P = 0.0282) all relative to the moderate progressing group as a reference. The lack of a strong genetic association is due to the small sample size, and that genetic variants relating to risk do not necessarily affect progression.

### Replication in an independent cohort

In order to ensure the generalizability and validity of the results, we replicated the subtype identification in an independent PDBP cohort. Details on differences between the training (PPMI) and replication (PDBP) cohorts can be found in the appropriately named **Online Results** section.

Figure 5 shows the identified subtypes in the independent PDBP cohort using the model developed on the PPMI dataset. We see that the identified subtypes in the PDBP cohort are similar to the ones in the PPMI dataset in terms of progression. Due to the limited length of the PDBP study (36 months), the visualization of progression space is shown through the 36 months follow-up from the baseline. The PPMI and PDPB cohorts are clinically different cohorts and recruited from different populations. The replication of our results in the PDBP cohort that was recruited with a different protocol shows the strength of our study’s methodology. We demonstrate that if we ascertain the same phenotypes using standardized scales, we can reliably discern the same subtypes and progression rates. This suggests that our results may be generalizable and the clinical subtypes reproducible.

**Fig 5.**
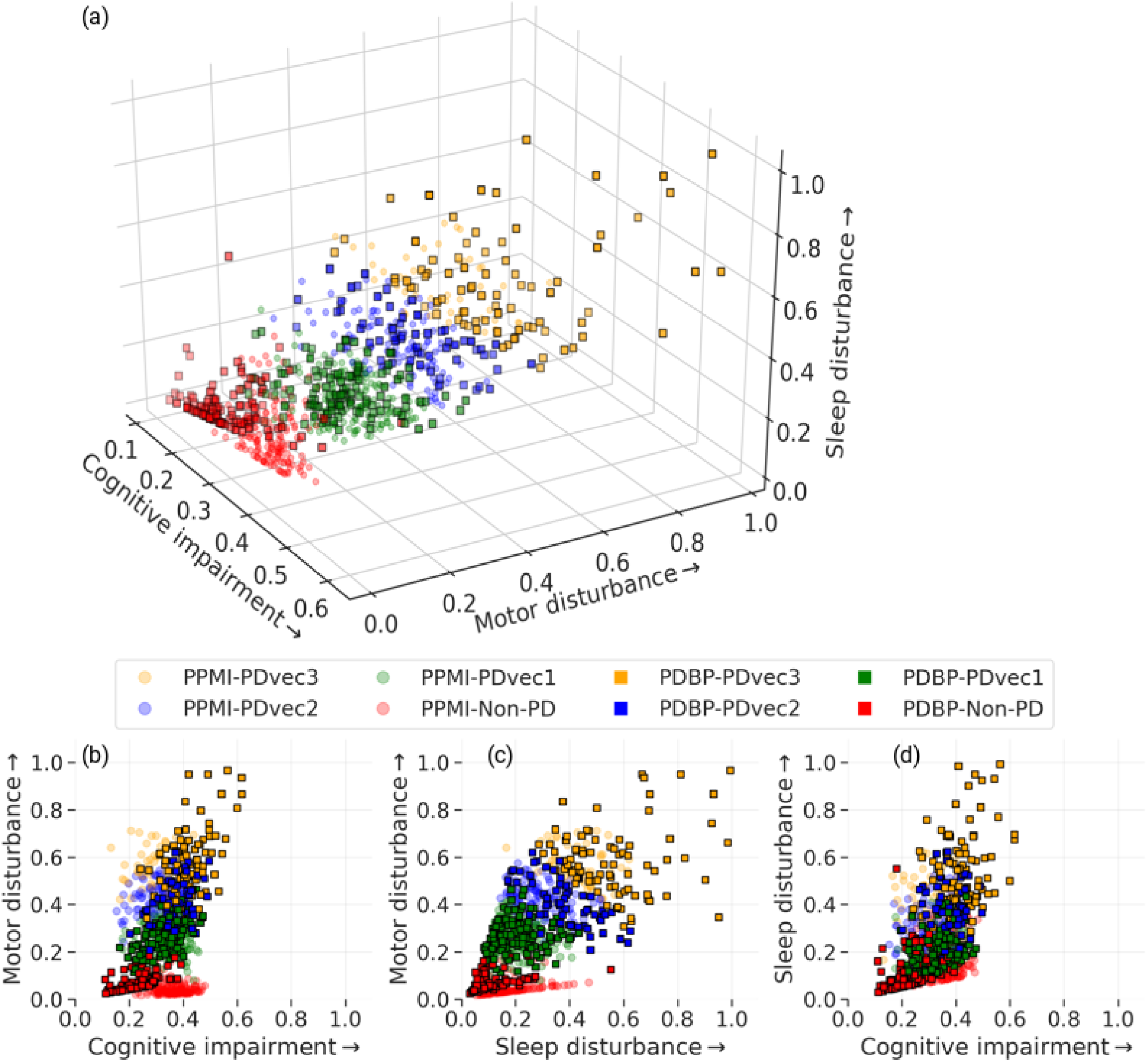
Shows the identified subtypes in the independent PDBP cohort using the model developed on the PPMI dataset. Similar PDBP and PPMI subtypes in terms of progression. (a) shows the view of all three dimensions, (b) view of the motor and cognitive dimensions, (b) view of motor and sleep dimensions, and (d) view of sleep and cognitive dimensions. The normalized progression space is shown through the 36 months follow up from baseline for both PPMI and PDBP datasets.

### Supervised early subtype prediction

Following the data-driven organization of subjects into progression subtypes and clustering them into three subtypes, we developed three models to predict patient progression class after 60 months based on varying input factors: (a) from baseline clinical factors, (b) from baseline and year one clinical factor, (c) biological and genetics measurements. **Figure 6a** and **Figure 6b** show the ROC (Receiver Operating Characteristic) curves of our multi-class supervised learning predictors. We correctly distinguish patients with PD based on baseline only input factors and predict their 60-month prognosis with an average AUC of 0.92 (95% CI: 0.94 ± 0.01 for PDvec1, 0.86 ± 0.01 for PDvec2, and 0.95 ± 0.02 for PDvec3) at cross-validation. The predictor built on baseline and year 1 data performs even better with an average AUC of 0.953 (95% CI: 0.97 ± 0.01 for PDvec1, 0.91 ± 0.02 for PDvec2, and 0.97 ± 0.01 for PDvec3) also at cross-validation. In **Figure 6d**, we have shown the PD subtype predictive performance at baseline, only using baseline data, and years after, as more data becomes available and combined with the baseline. The increased accuracy trend is due to the availability of more information about a subject. This approach is also practical in a clinical setting, as physicians will provide a better prognosis for patients after a one-year follow-up. Further details on feature importance contributing to the accuracy of these models can be found in the **Online Results** section entitled **Feature Importance**.

**Fig 6.**
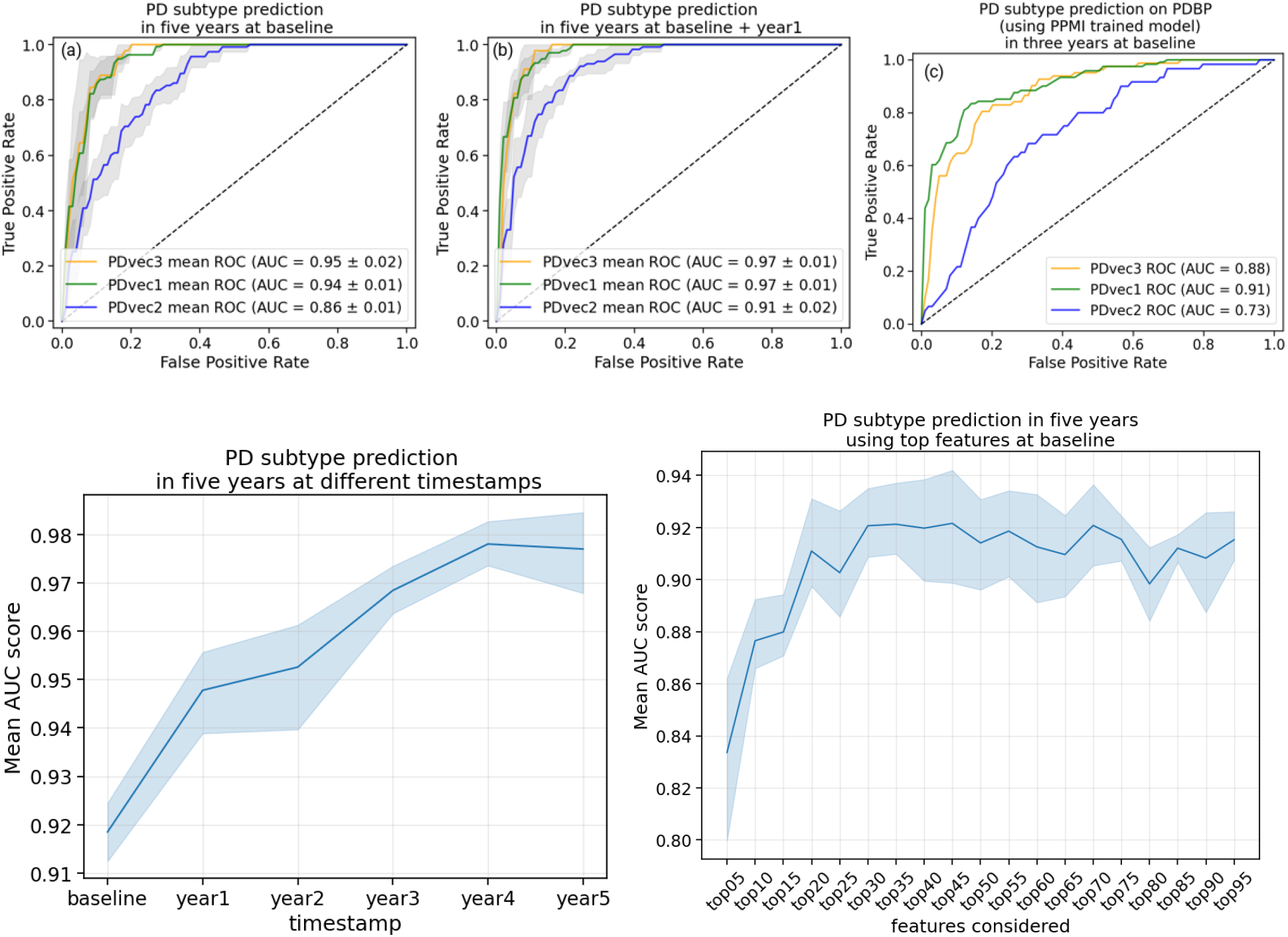
Shows the performance of Parkinson’s disease progression prediction models. (a) The ROC for the predictive model at baseline developed on the PPMI cohort evaluated using five-fold cross-validation. (b) The ROC for the predictive model developed on the baseline, and first-year data of the PPMI cohort evaluated using five-fold cross-validation. (c) The ROC for the predictive model developed on the PPMI baseline and tested on the PDBP cohort. (d) Performance of predictive models using data starting from baseline, only using baseline data, and years after, as more data becomes available and combined with the baseline. The y-axis shows the average AUC score across PD subtypes in the PPMI dataset. (e) Contribution of important features to achieve high accuracy. By including only 20 features, we can achieve an AUC of greater than 0.90.

Besides the cross-validation of predictive models in the PPMI cohort, we have also validated the accuracy of the predictive model in the independent PDBP cohort. The predictive model trained on the PPMI baseline data correctly distinguished PDBP patients with an AUC of 0.84 (ROC curves in **Figure 6c**). The replicated predictive model performs very well for PDvec1 and PDvec3 (AUC of 0.91 and 0.88, respectively). However, due to the small sample size, the predictive model does not predict as well on PDvec2 (AUC of 0.73). Fewer samples make up the PDvec2 cluster in the replication cohort, and it has been easier for the predictive model to predict the more extreme subtypes (i.e., PDvec1 and PDvec3). Despite the smaller sample size of the PDBP cohort, the results strongly validate our previous observations of distinct, computationally discernible subtypes within the PD population. This finding indicates that our methodology is robust, and our unique progression analysis and clustering approach result in the same clusters. In summary, we have mined data to identify three clinically-related constellations of symptoms naturally occurring within our longitudinal data that summarize PD progression (63.58%, 21.81%,14.61% variance loadings) comprised of factors relating to motor, sleep, and cognitive.

### Biomarker based prediction

There were 448 observations in total. We did not include plasma, CSF hemoglobin, and CSF glucosylceramide features because of their high missing data (>35%). Baseline CSF data were missing for p-tau in 51 patients, for total-tau in 26 patients, for abeta-42 in 20 patients, and alpha syn in 15 patients. Serum Nfl is missing in 22 participants, 17 participants did not have data for DNA GRS scores, and 36 participants had missing APOE status. The overall missing percentage for the above measurements was approximately 5%. We imputed the missing predictor variable data with means for numerical features and used most frequent category for categorical features. The remaining features include demographic information (education year, biological sex, birthdate, race), vital signs (weight, height, blood pressure), and family history (parents’ PD status) with no patients having missing data for them. For hg-38 genetic measurements, data was missing for 8 participants. We removed these patients from our study. Finally, we had 440 participants and 39 and 64 features for biomarkers and genetics measurements, respectively. All the genetic features are considered as categorical except DNA GRS. For the combined model, we concatenated both the biospecimen and genetic features. This concatenated vector was used as input for the classification of PD subtypes. The performance of PD progression prediction models using biomarkers and genetic measurements for the PPMI cohort is shown in **Figure 7** and **Table 1**. Additional notes on model interpretation as well as how the models deal with participants in the study that have changed diagnoses over time can be found in the **Online Results** in the appropriately named sections.

**Table 1.**
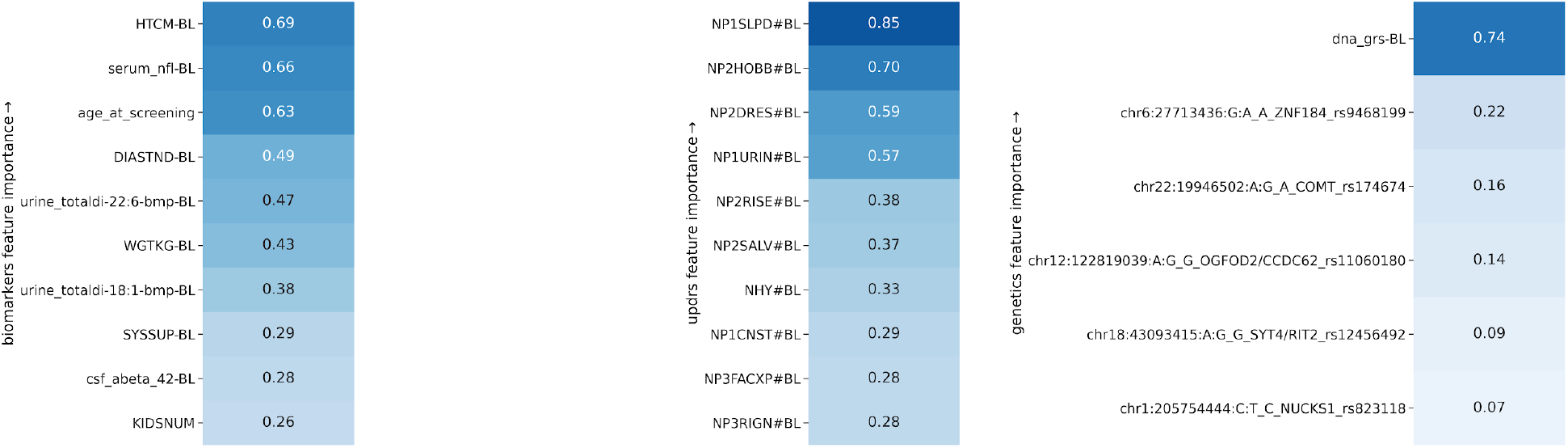
Table lists significant contributing clinical parameters (refer to **Table S1** for feature description) based on demographics, vital signs, baseline biospecimen, baseline MDS-UPDRS scores, and genetic measurements. The importance of each feature is relative.

**Fig 7.**
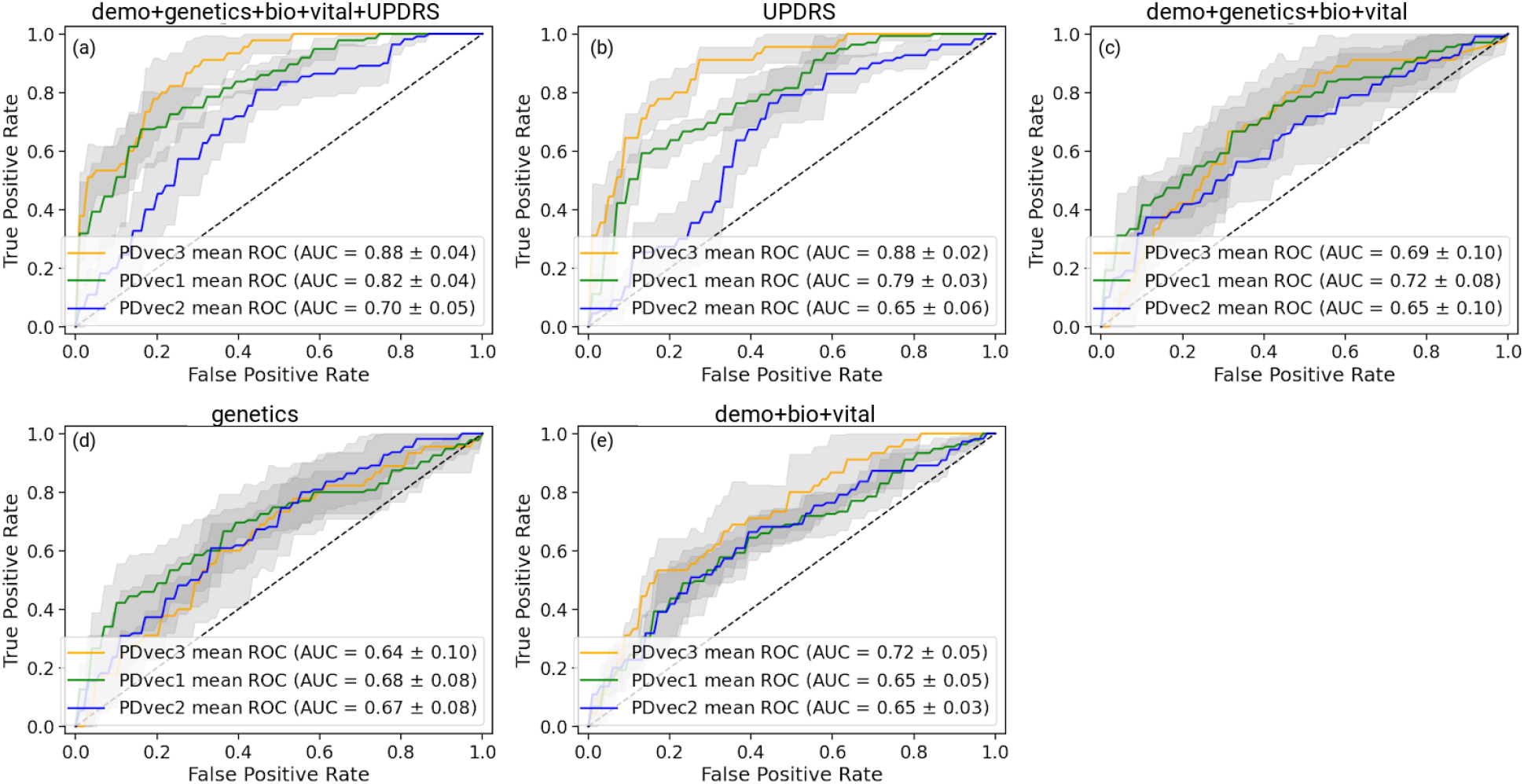
Shows the performance of Parkinson’s disease progression prediction models using biomarkers and genetic measurements for the PPMI cohort. All models are evaluated using five-fold cross-validation. From top left to bottom right: (a) The ROC for the predictive model using a combination of demographics (education, year, sex, race), biospecimen (cerebrospinal fluid, serum Nfl levels), genetics (hg genotype), vital signs (weight, height, blood pressure) and UPDRS measurements. (b) The ROC for the predictive model developed on UPDRS scores. (c) The ROC for the predictive model developed using demographics, genetics, vital signs, and biospecimen measurements. (d) The ROC for the predictive model developed on genetic measurements (e) The ROC for the predictive model uses only demographics, vital signs, and biospecimen measurements.

Biomarkers, such as age, height, weight, and CSF measurements, are shown to be essential features in predicting the subtypes at baseline (shown in **Table 1**). The mean AUC score is 0.67 (**Figure 7e)** using biospecimen, vital signs, and demographics. In comparison, genetic features show slightly lower performance (AUC score 0.66, **Figure 7d**). A combination of demographics, biospecimen, vital signs, genetics, and UPDRS is the best performing model (AUC score 0.80, **Figure 7a**).

## Discussion

Prediction of disease and disease course is a critical challenge in the patient counseling, care, treatment, and research of complex heterogeneous diseases. Within PD, meeting this challenge would allow appropriate planning for patients and symptom-specific care (for example mitigating the chance of falls, identifying patients at high risk for cognitive decline or rapid progression, etc.). Perhaps even more importantly at this time, prediction tools would facilitate more efficient execution of clinical trials. With models predicting a patient-specific disease course, clinical trials could be shorter, smaller, and would be more likely to detect smaller effects, thus, decreasing the cost of phase 3 trials dramatically and essentially reducing the exposure of pharmaceutical companies to a typically expensive and failure-prone area.

We previously had considerable success in constructing, validating, and replicating a model that allows a data-driven diagnosis of PD and the differentiation of PD-mimic disorders, such as those patients who have parkinsonism without evidence of dopaminergic dysfunction (Nalls et al. 2015). We set out to expand this work by attempting to use a novel approach to 1) define natural subtypes of the disease, 2) attempt to predict these subtypes at baseline, and 3) identify progression rates within each subtype and project progression velocity.

While the work here represents a step forward in our efforts to sub-categorize and predict PD, much more needs to be done. The application of data-driven efforts to complex problems such as this is encouraging; however, the primary limitation of such approaches is that they require large datasets to facilitate model construction, validation, and replication. These datasets should include standardized phenotype collection and recording to achieve the most powerful predictions. Collecting such data is a challenge in PD, with relatively few cohorts available with deep, wide, well-curated data. Thus, a critical need is the expansion or replication of efforts such as PPMI or PDBP, importantly with a model that allows unfettered access to the associated data; the cost associated with this type of data collection is large, but these are an essential resource in our efforts in PD research.

A recent study used cluster analysis to identify patient subtypes and their corresponding progression rates (Fereshtehnejad et al. 2017), although these used percentile cutoffs and are not completely data-driven in nature. However, this study evaluated clusters according to only two-time points, baseline, and short-term follow-up, that were aggregated into a Global Composite Outcome score. In return, the subtypes did not capture the fluctuations in the prognosis of subtypes. More recently, a study (Zhang et al. 2019) used a Long-Short Term Memory-based deep learning algorithm to discover PD subtypes, with each subtype showing different progression rates. The loss of interpretability with deep learning models makes their approach less suitable for practical purposes. Another study (Krishnagopal et al. 2019) proposed a trajectory-based clustering algorithm to create patient clusters based on trajectory similarity. The algorithm gives equal importance to all the features; however, PD is a multi-dimensional spectrum of symptoms with overlapping features derived from simultaneous pathological processes (Marras and Chaudhuri 2016). Finally, in order to be used in practice, subtyping solutions need to be replicated in a different cohort to show the reliability of methods in assigning individual patients to a subtype. Additionally, none of these previous studies used completely independent replication data.

Our findings can also have implications for the day-to-day practice of clinicians. Movement disorders specialists often use screening tools such as MDS-UPDRS to assess a patient’s progression and response to treatment. However, performing these clinical assessments requires experience, expertise, and time, which hinders its widespread use by other clinicians (and even neurologists who are not trained in movement disorders). Underutilization of clinical assessment tools can lead to the suboptimal characterization of PD patients and their clinical course, which in turn impacts their care. Our study is one of the first of its kind which systematically assessed the accuracy of each feature of MDS-UPDRS in predicting PD’s course. For example, daytime sleepiness (NP1SLPD) was found to have the highest importance in clinical progression, followed by doing hobbies and activities (NP2HOBB), dressing (NP2DRES), and urinary problems (NP1URIN). Knowing the clinical features with the highest yield in course prediction can help clinicians to tailor their assessment and better inform patients about their disease course. In addition, shortened versions of comprehensive assessment tools can be utilized to address specific clinical questions. Surprisingly, none of the genetic markers studied had high accuracy in clinical course prediction. Our work initiates multiple questions that are worth exploring in the future. The progression space seems to stabilize after three years from the baseline. It will be interesting to predict how much time (from baseline) is required to provide reliable predictions about the PD subtypes. Secondly, fast progressors do not worsen with the multiple symptoms such as Epworth and MDS-UPDRS scores (**Figure S5b**) from the fourth to the fifth year, while other subtypes do. It raises the question of whether the fast progressors reach the saturation point after some time from baseline. It will be useful if we can look for similar patterns in other PD datasets. Incorporating imaging data for PD subtypes is also an exciting direction to pursue in the future. Finally, dramatic increases in Nfl and high baseline levels of Nfl could be an indicator of potential rapid progression.

## Conclusions

In this study, we addressed the complexities of PD. We integrated unlabeled, multimodal, and longitudinal data. The longitudinal data had a long-term nature, and we were interested in capturing the overall pattern of the individual’s trajectories. Vectorization and NMF methods were the most successful approaches for extracting long-term trajectories. Using comprehensive multi-modal data helped us to develop an embedded space. This space was crucial for understanding the trajectories and dimensions in which the individuals traverse. Having this easily interpretable space, we were able to use a GMM unsupervised learning approach to identify new subtypes of the disorder based on disease progression. We also provided an in-depth analysis of these subtypes. Furthermore, we developed predictive models for early diagnosis, prognosis, and clinical trial stratification.

This work provides data-driven subtypes in distinct progression stages of PD and discusses an approach to predict the future rate of progression years from baseline using longitudinal clinical data. Predicting disease progression is a paramount challenge in treating and curing several complex diseases. This study is a step forward toward designing sophisticated machine-learning paradigms to facilitate the early diagnosis of PD progression and longitudinal biomarker discovery such as our finding of elevated Nfl in fast progressors (both at baseline and with regard to the rate of change per year). Predicting PD progression rates would lead to better patient-specific attention by recognizing the patients with a swift rate of progression at an early stage. The proposed disease progression and trajectory prediction algorithms can help healthcare providers to develop a methodical and organized course for clinical tests, which can be much more concise and effective in detection. These adaptations and modifications in clinics may help to diminish treatment and therapy costs for PD. Further, the capability to anticipate the trajectory of impending PD progression at the early stages of the disease is an advancement toward uncovering novel treatments for PD modification. The proposed analysis provides insights to inhibit or decelerate the progression of PD-related symptoms and subsequent deterioration in the characteristics of life that are accompanied by the disease.

## Acknowledgments

We thank the patients and their families who contributed to this research. This work was supported in part by the Intramural Research Program of the National Institute on Aging and the National Institute of Neurological Disorders and Stroke, National Institutes of Health, Department of Health and Human Services (project _ZO1 AG000949, ZIA-NS003154), and the Michael J Fox Foundation. This work has also been supported in part through the grant 1U54GM114838 awarded by NIGMS through funds provided by the trans-NIH big data to Knowledge (BD2K) initiative (www.bd2k.nih.gov). Data used in the preparation of this article were obtained from the Parkinson’s Progression Markers Initiative (PPMI) database (www.ppmi-info.org/data). For up-to-date information on the study, visit www.ppmi-info.org. PPMI – a public-private partnership – is funded by the Michael J. Fox Foundation for Parkinson’s Research and funding partners, including Abbvie, Avid Radiopharmaceuticals, Biogen Idec, Bristol-Myers Squibb, Covance, Eli Lilly & Co., F. Hoffman-La Roche, Ltd., GE Healthcare, Genentech, GlaxoSmithKline, Lundbeck, Merck, MesoScale, Piramal, Pfizer, and UCB. Data and biospecimens used in the preparation of this manuscript were obtained from the Parkinson’s Disease Biomarkers Program (PDBP) Consortium, part of the National Institute of Neurological Disorders and Stroke at the National Institutes of Health. Investigators include: Roger Albin, Roy Alcalay, Alberto Ascherio, DuBois Bowman, Alice Chen-Plotkin, Ted Dawson, Richard Dewey, Dwight German, Xuemei Huang, Rachel Saunders-Pullman, Liana Rosenthal, Clemens Scherzer, David Vaillancourt, Vladislav Petyuk, Andy West and Jing Zhang. The PDBP Investigators have not participated in reviewing the data analysis or content of the manuscript.

