## Supplementary Information for "Identification and prediction of Parkinson’s disease subtypes and progression using machine learning in two cohorts"

### Supplementary materials

#### Online methods

##### Data preprocessing

As the clinical features have varying directionalities, features were transformed to ensure the highest values uniformly represent the worst outcome while the lower value corresponds to greater health. To identify the features not following this pattern, we conducted a two-sample one-tailed t-test (null hypothesis: μ HC >= μ PD ) with cases versus controls as two samples. We transformed those features that showed a *p*-value of less than 0.05 (5% significance test). We further verified the transformations by manually reviewing the distribution of each feature.

We performed feature clipping to minimize the influence of extreme outliers in the data. We limited all the features values between the range given by the second and 98$^{th}$ percentiles.
Only a few features had residual missingness that was distributed randomly across the patients at a rate of <5%. For these features, we performed data imputation using linear interpolation longitudinally (i.e., across visits) for each feature.
Then, we transformed the dataset into a mathematically meaningful and naturally interpretable format. To achieve this objective, we a) *vectorized* and b) *normalized* all longitudinal data. Specifically, we first vectorized by transforming all observations of a particular parameter in a column vector, then appended all parameters together. We then used the *min-max* method to normalize the data.

##### NMF

Mathematically, NMF factorizes (deconstructs) the data into two matrices. Given a non-negative matrix $X \in R^{m\times n}$, a non-negative decomposition of the matrix $X$ is a pair of non-negative matrices $U \in R^{m\times p}$ and $V \in R^{p\times n}$such that $X = UV$. A large number of patient parameters are aggregated in a model that represents the underlying progression concept. In this particular use case of NMF, the matrix $U$ contains the progression space latent vectors, and the second matrix $V$ contains progression stand indicators corresponding to the latent vectors. Latent variables link observation data in the real world to symbolic data in the modeled world. By further looking into the matrix with progression space's latent vectors, we can identify the mapping and, consequently, the implications (symbolic dimensions of the modeled progression space).

#### Latent Space Adjustment

The progression of space latent vectors (matrix $U$) learned by NMF shows some weight sharing among the projected dimensions. This weight sharing can be attributed to the presence of correlation in the data between symptomatologies. To represent the progression space so that each progression indicator shows progression velocity for each symptom, we need to adjust the progression space. We performed transformation on NMF learned progression space by taking the weighted sum of the progression indicators. These weights represent the contribution of each dimension for distinct symptomatologies.

$$i^{symptom\ s}_{new} = \Sigma_d c^{symptom\ s}_{dimension\ d} * i^{dimension\ d}_{old}$$

Here, $c^{symptom\ type}_{dimension\ (k)}$ denotes the contribution of dimension k for features belonging to symptom $s$, $i^{dimension\ d}_{old}$ is the latent indicator learned by NMF, and $i^{symptom\ s}_{new}$ is the new indicator after the adjustment. The component contribution is calculated using the learned $U$ matrix.

Through our use of NMF, we identified progressive features based on motor, cognitive, and sleep-based disturbances. Following this, unsupervised learning via Gaussian Mixture Models (GMM) (McLachlan and Basford 1988) allowed the data to naturally self-organize into different groups relating to velocity of decline across these three categories, from non-PD controls representing normal aging to PD subtypes. GMM is a variant of mixture models, compared to other methods, the parametrization of a GMM allows it to efficiently capture products of variations in natural phenomena where the data is assumed generated from an independent and identically distributed (i.i.d.) mixture of Gaussian (normal) distributions. The assumption of normal distribution (and therefore, the use of GMM) is often used for population-based cohort phenomenon (Prentice 1986).

We use the Bayesian Information Criterion (BIC) to select the number of PD clusters (subtypes) (Schwarz 1978). The BIC method recovers the true number of components in the asymptotic regime (i.e., much data is available, and we assume that the data was generated i.i.d. from a mixture of Gaussian distributions). To replicate the subtype identification, we applied the GMM model developed in the PPMI data to an independent cohort with varying recruitment strategy and design: the PDBP cohort.

#### Model Evaluation and Tuning

**Algorithm 1:** Model Evaluation Procedure

```
for each iteration i=1,2,..I do
    Divide the dataset into train (80%) and test data (20%) at random;
    Divide the train into K cross-validation subfolds at random;
    for each fold k=1,2,..K // CV loop do
        validation = fold k;
        subtrain = all folds other than k;
        Train model with every hyperparameter on subtrain;
        Evaluate it on validation;
    end
    Calculate the average metrics score on validation over the K folds
      for every hyperparameter;
    Choose the best hyperparameter setting;
    Train a model with the best hyperparameter on train;
    Evaluate its performance on test;
end
Calculate the mean accuracy over all I iterations on test data;
```

Fig S1. The workflow of predictive modeling evaluation and hyper-parameter tuning.

Table S1. Shows the description of PPMI clinical assessment features and their labels.

| | |
|---|---|
| **MDS-UPDRS Part 1** | |
| *NP1DPRS* | Depressed mood |
| *NP1ANXS* | Anxious mood |
| *NP1SLPN* | Sleep problems |
| *NP1PAIN* | Pain and other sensations |
| *NP1CNST* | Constipation problems |
| *NP1FATG* | Fatigue |
| *NP1DDS* | Dopamine dysregulation syndrome |
| *NP1SLPD* | Daytime sleepiness |
| *NP1URIN* | Urinary problems |
| *NP1HALL* | Hallucinations |
| *NP1APAT* | Apathy |
| *NP1COG* | Cognitive impairment |
| *NP1LTHD* | Lightheadedness on standing |
| **MDS-UPDRS Part 2** | |
| *NP2SALV* | Saliva and drooling |
| *NP2EAT* | Eating tasks |
| *NP2DRES* | Dressing |
| *NP2HYGN* | Hygiene |
| *NP2HWRT* | Handwriting |
| *NP2HOBB* | Doing hobbies and other activities |
| *NP2RISE* | Getting out of bed, car, deep chair |
| *NP2FREZ* | Freezing |
| *NP2SWAL* | Chewing and swallowing |
| *NP2TURN* | Turning in bed |
| *NP2WALK* | Walking and balance |
| *NP2SPCH* | Speech |
| *NP2TRMR* | Tremor |
| **MDS-UPDRS Part 3** | |
| *NP3SPCH* | Speech |
| *NP3FACXP* | Facial expression |
| *NP3RIGN* | Rigidity – neck |
| *NP3RIGRU* | Rigidity – RUE |
| *NP3RIGRL* | Rigidity – RLE |
| *NP3RIGLL* | Rigidity – LLE |
| *NP3FTAPR* | Finger tapping right hand |
| *NP3FTAPL* | Finger tapping left hand |
| *NP3HMOVL* | Hand movements – left hand |
| *NP3HMOVR* | Hand movements – right hand |
| *NP3PRSPR* | Pronation-supination – right hand |
| *NP3PRSPL* | Pronation-supination – left hand |
| *NP3TTAPL* | Toe tapping – left foot |
| *NP3TTAPR* | Toe tapping – right foot |
| *NP3LGAGR* | Leg agility – right leg |
| *NP3LGAGL* | Leg agility – left leg |

| | |
|---|---|
| *NP3RISNG* | Arising from chair |
| *NP3POSTR* | Posture |
| *NP3BRADY* | Global Spontaneity of movement |
| *NP3KTRML* | Kinetic tremor – left hand |
| *NP3RTARL* | Rest tremor amplitude – RLE |
| *NP3RTALL* | Rest tremor amplitude – LLE |
| *NHY* | Hoehn and Yahr stage |
| *DYSKPRES* | Presence of dyskinesias |
| *NP3FRZGT* | Freezing of gait |
| *NP3RTALJ* | Rest tremor amplitude - Lip/jaw |
| *NP3RTCON* | Constancy of rest |
| *NP3GAIT* | Gait |
| *NP3RTALU* | Rest tremor amplitude - LUE |
| *NP3RTARU* | Rest tremor amplitude - RUE |
| *NP3PTRMR* | Pronation - Supination Movements - Right Hand |
| *NP3PTRML* | Pronation - Supination Movements - Left Hand |
| *NP3PSTBL* | Postural stability |
| **MoCA** | |
| *Naming* | Naming total score |
| *Language* | Language total score |
| *Delayed recall* | Delayed recall total score |
| *visuospatial* | Visuospatial total score |
| *attention* | Attention total score |
| *MCAVFNUM* | Verbal fluency |
| *MCAABSTR* | Abstraction |
| *MCATOT* | MoCA total score |
| **HVLT** | |
| *HVLTRT1* | Immediate Recall Trial 1 |
| *HVLTRT3* | Immediate Recall Trial 3 |
| *HVLTRT2* | Immediate Recall Trial 2 |
| *HVLTRDLY* | Delayed Recall |
| *HVLTREC* | Recognition |
| *HVLTFPRL* | Recognition – false positives, related |
| **LNS** | |
| *LNS_TOTRAW* | LNS-Sum questions 1-7 |
| **QUIP** | |
| *TMSEX* | Think having too much sex behavior |
| *TMGAMBLE* | Think having too much gambling behavior |
| *TMTORACT* | Too much time on recreational activities |
| *CNTRLBUY* | Difficulty controlling your buy behaviors |
| *TMTMTACT* | Too much time on motor activities |
| *TMTRWD* | Too much time on walking/driving activities |
| *TMEAT* | Think having too much eating behavior |
| *CNTRLGMB* | Difficulty controlling your gambling behaviors |
| *CNTRLSEX* | Difficulty controlling your sex behaviors |
| *TMBUY* | Think having too much buying behavior |

| | |
|---|---|
| *CNTRLEAT* | Difficulty controlling your eat behaviors |
| **RBDSQ** | |
| *DRMAGRAC* | Dreams frequently have aggressive or action-packed content |
| *SLPINJUR* | I (almost) hurt my bed partner or myself |
| *DRMVERBL* | In my dreams: speaking, shouting, swearing |
| *DRMUMV* | In my dreams: gestures, complex movements useless during sleep |
| *DRMOBJFL* | In my dreams: things fell down around the bed |
| *MVAWAKEN* | It happens that my movements awake me |
| *SLPDSTRB* | My sleep is frequently disturbed |
| *SLPLMBMV* | Know my arms and legs move when asleep |
| *RLS* | had RLS |
| *STROKE* | Disease of nervous system: stroke |
| *DEPRS* | Disease of nervous system: depression |
| *NARCLPSY* | had narcolepsy |
| *DRMREMEM* | remember the content of my dreams well |
| *DRMNOCTB* | dream contents mostly match my nocturnal behaviour |
| *BRNINFM* | had inflammatory disease of the brain |
| *DRMVIVID* | sometimes have very vivid dreams |
| *HETRA* | had head trauma |
| *EPILEPSY* | had epilepsy |
| *DRMFIGHT* | In my dreams: sudden limb movements |
| **EPS** | |
| *ESS2* | Doze off or fall asleep while watching TV |
| *ESS1* | Sitting and reading |
| *ESS3* | Sitting, inactive in a public place |
| *ESS4* | As a passenger in a car for an hour without a break |
| *ESS5* | Lying down to rest in the afternoon |
| *ESS6* | Sitting and talking to someone |
| *ESS7* | Sitting quietly after a lunch |
| *ESS8* | In a car, while stopped for a few minutes |
| **SCOPA-AUT** | |
| *Gastrointestinal upper* | Had difficulty swallowing or have choked + Has saliva dribbled out of your mouth + Has food become stuck in your throat |
| *Gastrointestinal lower* | Have feeling during meal that you were full very quickly + Had problems with constipation + Had to strain hard to press stools + Had involuntary loss of stools |
| *thermoregulatory* | trouble tolerating cold + trouble tolerating hot |
| *pupillomotor* | eyes ever been over-sensitive to bright light |
| *skin* | perspired excessively during the day + during the night |
| *cardiovascular* | feeling of either becoming light-headed + light-headed after standing for some time + fainted in the past 6 months |
| *urinary* | difficulty retaining urine + involuntary loss of urine + stream of urine been weak + pass urine at night + urine your bladder was not completely empty + urine again within 2 hours of the previous time |
| **Semantic Fluency** | |
| *VLTANIM* | Total number of animals |
| *VLTVEG* | Total number of vegetables |

| | |
|---|---|
| *VLTFRUIT* | Total number of fruits |
| **STAI** | |
| *a_state* | Anxiety state score |
| *a_trait* | Anxiety trait score |
| **BENTON** | |
| *JLO_TOTRAW* | judgment line of action total raw score |
| **Geriatric** | |
| *total* | geriatric depression total score |
| **Neuro Cranial** | |
| *CN346RSP, CN8RSP, CN7RSP, CN2RSP, CN910RSP, CN12RSP, CN5RSP, CN11RSP* | abnormality in Cranial Nerves |
| **SDM** | |
| *SDMTOTAL* | total symbol digit modalities test |

### Online results

#### Interpretation of the Latent Space

To understand the interpretation of PD progressions space dimensions, **Figure S2** shows the mapping guide for how the PPMI's high-dimensional space of 138 different clinical parameters is mapped to the three-dimensional embedding of Parkinson's disease progression space. The features are grouped together to represent coherent skills. The leftmost component in **Figure S2** mainly constitutes the questionnaire associated with sleep and mood problems, such as dream, fatigue, anxiety, and depression. The middle component represents questions related to motor skills such as speech, facial expression, tremor, and rigidity. The third component represents questions related to cognitive skills, such as cognitive assessment and verbal learning tests. Therefore, the columns represent the projected three dimensions, i.e., motor, cognitive, and sleep-related trajectories, and the rows are the PPMI clinical parameters. This interpretable mapping is due to the property of NMF to group features showing similar variations in the data. This figure allows us to not only observe the conversion but also the heterogeneity of some clinical parameters, for instance how some of the Epworth Sleepiness Scale parameters reflect both sleep and cognitive disorders, and some reflect both sleep and movement disorders. We also looked at the features that seem to be incorrectly assigned, such as cognition (NP1COG) in the motor, and neurocranial (CN346RSP) in sleep. We find that the responses to these questions show minimal variation across subjects, which might make NMF assign them to any of the components. In comparisons of the eigenvalues within the NMF decomposition, the projected motor dimension was responsible for 63.58% of the explained variance, followed by the sleep dimension (21.81·%), and cognitive dimension (14.61%).

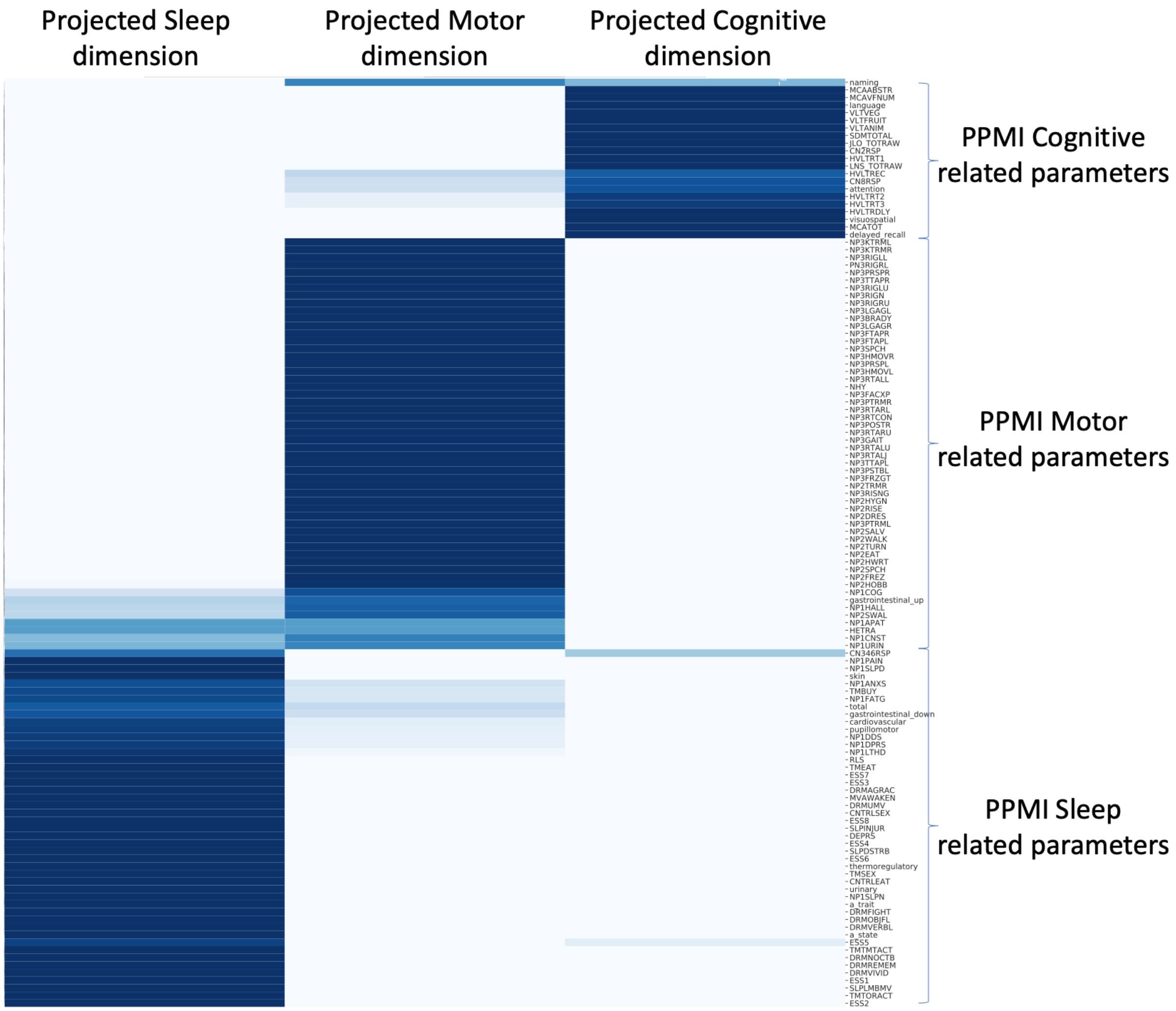

Fig S2. Shows how each 122 different input parameters have been projected into the new dimension of the Parkinson's progression space (cognitive, motor, and sleep dimensions). Darker colors represent strong mapping. The mapping is shown at the final visit after z-score normalization; it is similar for other visits as well (Figure not shown).

#### Five Year Progression Space

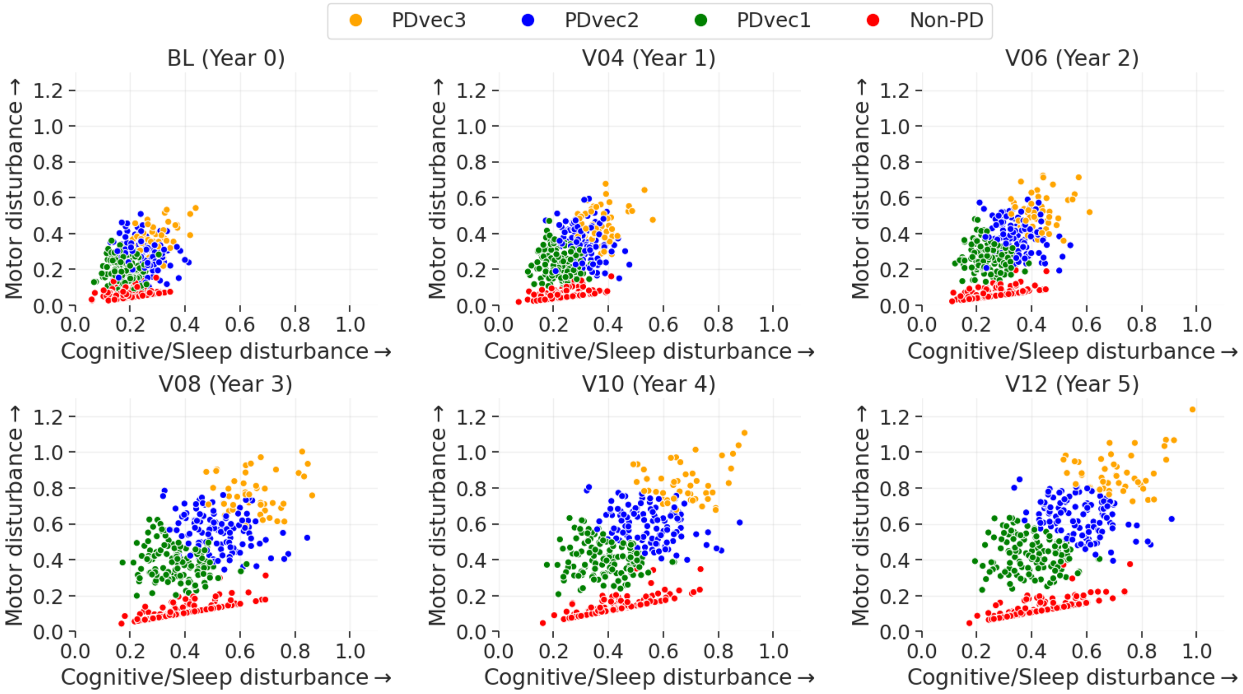

Fig S3. Visualization of two-dimensional progression space of PD subtypes at the end of every year, showing their normalized trajectory. (BL-baseline, V04-Year1, V06-Year2, V08-Year3, V10-Year4, V12-Year5)

**Figure S3** shows the disease trajectory of different PD subtypes. The progression space shows the gradual and linear change for all the subjects. Furthermore, the progression space tends to stabilize at the end of the third year. In this way, our model can capture patients' nuanced behavior showing their progression along with different skills. It is interesting to observe that a significant decline occurs between the second and third year for the subjects in our analysis. In terms of characteristics of PDs identified subtypes, **Figure S4** demonstrated how cognitive, motor and sleep-related symptoms differ within each PDs subtype and in controls. There is a clear trend for increased cognitive, sleep, and motor disturbances after five years in fast progressors compared to the slower progressing subtypes. The slowest progressive subtypes (PDvec1) show a mild decline for motor dimension but less change for sleep and cognitive dimensions. We can observe that the difference in progression rates between controls and fastest progressive subtypes is mainly along the motor dimension followed by sleep and then the cognitive dimension.

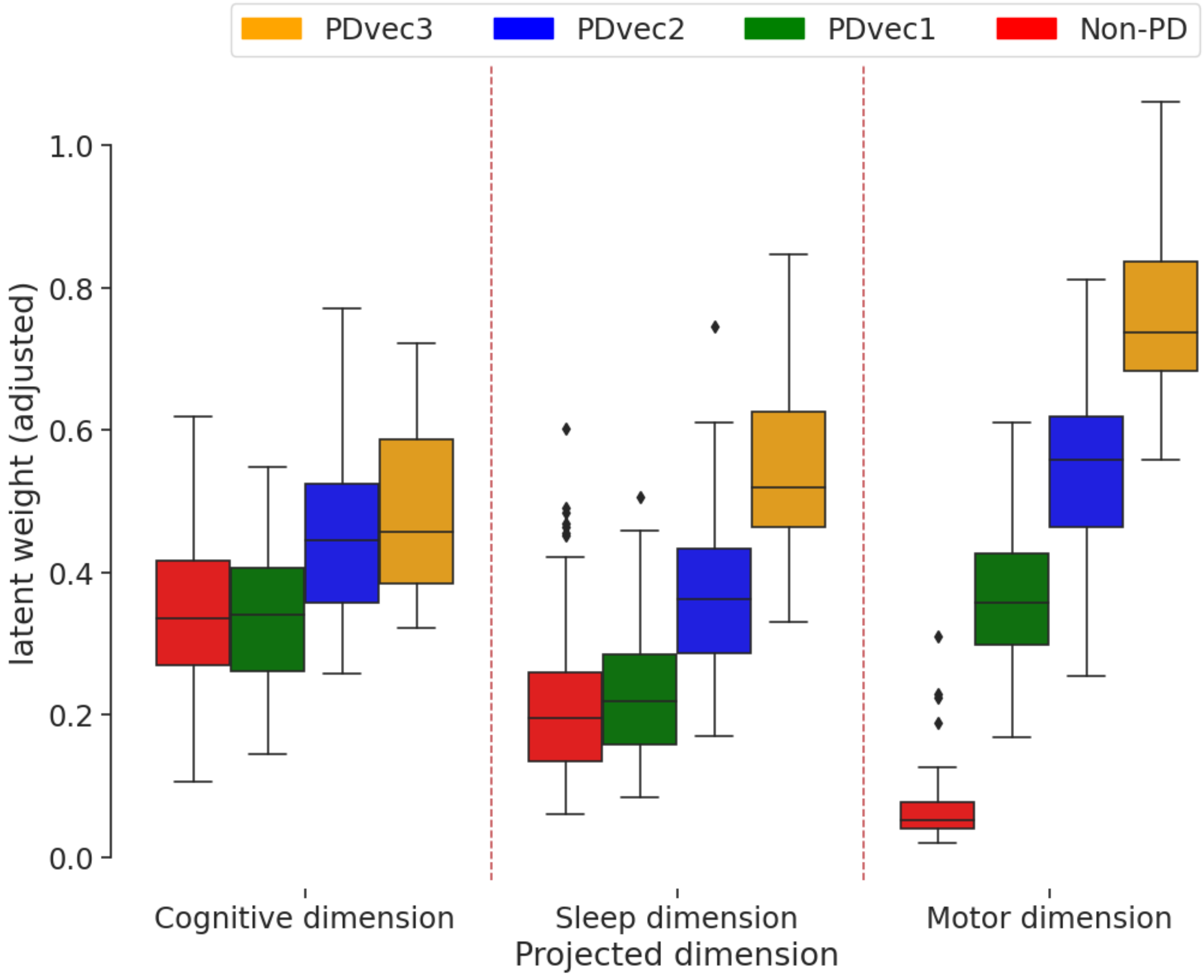

Fig S4. Shows the distribution of projected dimensions (cognitive, motor, and sleep) weights for each Parkinson's category and healthy control after five years. An increase in values along either direction reflects the increase in the disturbance. PDvec3 has the highest motor and sleep disturbance, as well as the highest cognitive impairment.

**Figure S5** shows the progression of each PD subtype overtime at baseline and after 12 months, 24, 36, 48 months, and 60 months. To better understand the clinical presentation of the three identified subtypes, **Figure S5(a) and S5(b)** demonstrates the three main projected dimensions (motor, cognitive, and sleep-related disturbances), as well as actual clinical values of each subtype overtime for UPDRS-Part I, Part II, Part III, as well as Hopkins Verbal Learning Test, Symbol Digit Modalities Test, Semantic Fluency test, Epworth Sleepiness Scale, State-Trait Anxiety Inventory for Adults, and Geriatric Depression Scale.

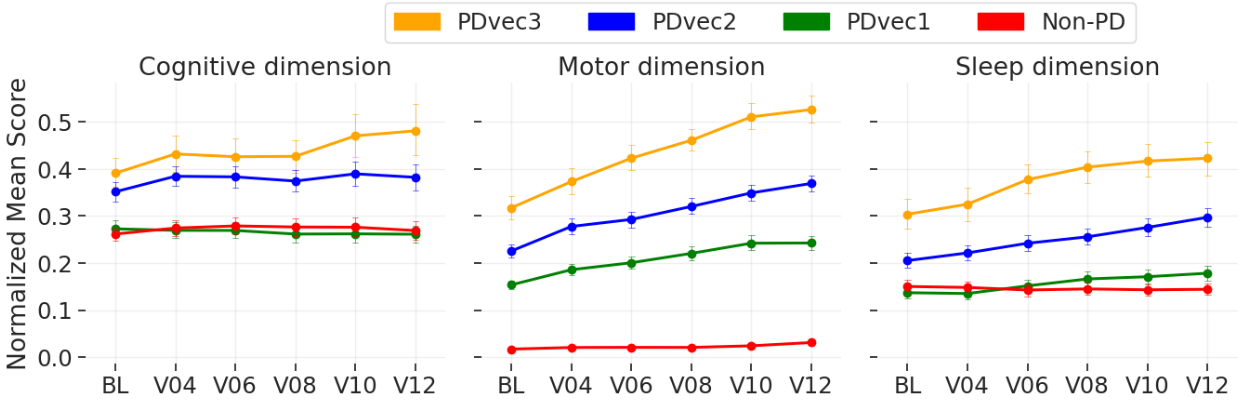

Fig S5(a) shows the progression of each PD subtype over time for motor, sleep, and cognitive dimension overtime on the preprocessed values.

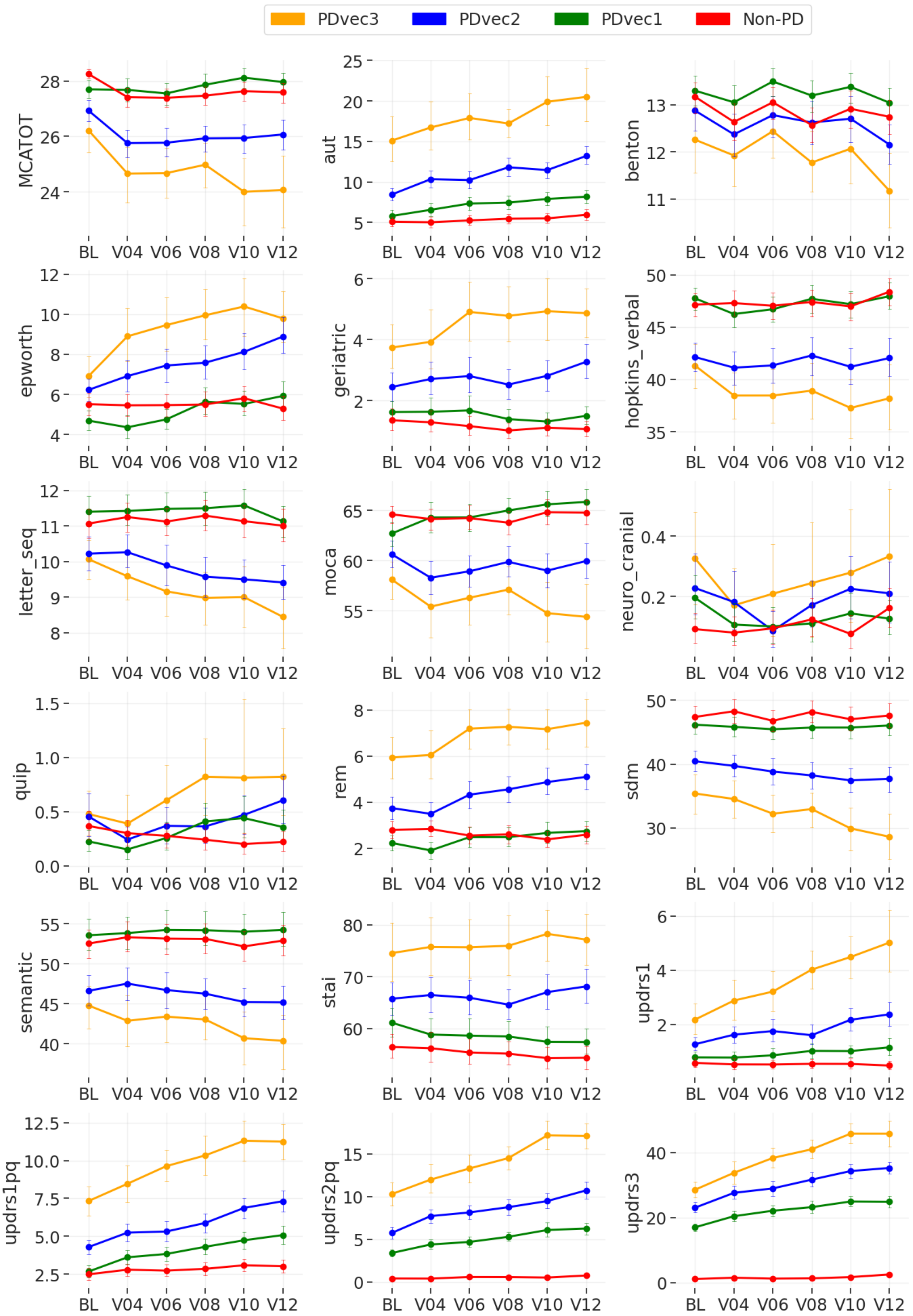

PDvec3
PDvec2
PDvec1
Non-PD
MCATOT
aut
benton
epworth
geriatric
hopkins_verbal
letter_seq
moca
neuro_cranial
quip
rem
sdm
semantic
stai
updrs1
updrs1pq
updrs2pq
updrs3
BL V04 V06 V08 V10 V12

Fig S5(b). Shows the progression of each PD subtype over time. The graphs demonstrate the actual clinical values of each subtype overtime for UPDRS-Part I, Part 2, Part 3, as well as Hopkins Verbal Learning Test, Symbol Digit Modalities Test, Semantic Fluency test, Epworth Sleepiness Scale, State-Trait Anxiety Inventory for Adults, and Geriatric Depression Scale. BL: Baseline. V04: visit number 4 after 12 months. V06: visit number 6 after 24 months. V08: visit number 8 after 36 months. V10: visit number 10 after 48 months. V12: visit number 12 after 60 months.

#### Additional Details on Replication Cohort

**Figure S6** shows PPMI and PDBP cohorts are similarly distributed; hence, they are suitable for replication and validation. Furthermore, we have performed the two-sample t-test for quantified replication cohort validation analysis (Table 1).

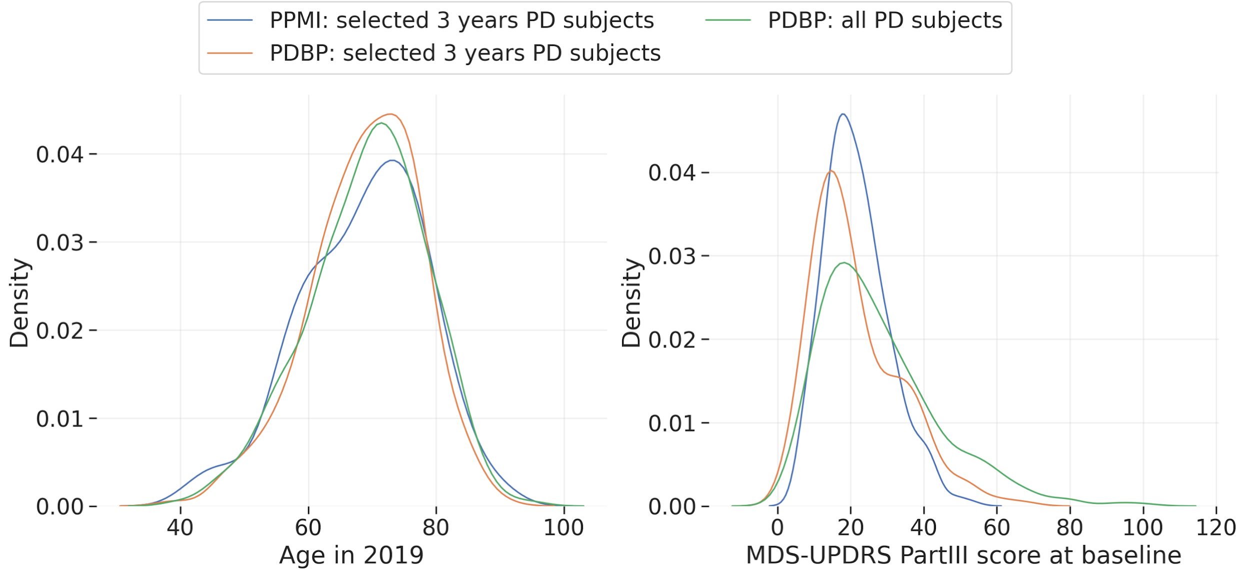

Fig S6. Kernel Density Estimation (KDE) analysis of Age and MDS-UPDRS Part III (objective motor symptom examination by a trained neurologist) in PPMI and PDBP cohorts. (a) shows the density of Parkinson's participant's age in the 3-years PPMI, PDBP, and 3-years PDBP datasets, and (b) shows the distribution of Parkinson's participant's MDS-UPDRS Part III at baseline in the 3-years PPMI, PDBP, and 3-years PDBP datasets. The three density functions in both figures are similar showing the validity of statistical replication.

| **PPMI vs PDBP (after 3 years)** | **t-value (95% CI)** | **p-value (95% CI)** |
|---|---|---|
| Age in 2019 | -0.41 | 0.68 |
| MDS UPDRS PartIII | 0.29 | 0.77 |

Table S2: Two-sample t-test for quantified replication cohort validation analysis. PPMI vs. PDBP (selected participants with 3 years of data).

#### Feature Importance

The predictive model was also analyzed to identify the feature importance in predicting PD subtypes. Feature importance is determined by calculating the relative influence of each variable, which is typically given by information gain/entropy, and how much the variable contributes to the accuracy (**Figure 6e**). We further scaled each feature's importance between 0 and 1. **Table S3** shows the top-50 features identified by our predictive model.

Table S3(a). Shows the summary of clinical parameters (top 50 features) to the prediction models ordered by their importance. The value indicates the scaled importance of the variables in predicting the PD subtypes. Table lists significantly contributing clinical parameters based on the baseline model and on model using both baseline and year 1 data.

| clinical feature importance (Model: baseline) → | baseline |
|---|---|
| NHY | 1.00 |
| NP3BRADY | 0.33 |
| NP3FACXP | 0.26 |
| NP2TRMR | 0.24 |
| urinary | 0.12 |
| SDMTOTAL | 0.09 |
| VLTANIM | 0.07 |
| NP1SLPD | 0.07 |
| NP2HOBB | 0.06 |
| gastrointestinal_down | 0.06 |
| NP3FTAPR | 0.05 |
| NP3RIGN | 0.05 |
| HVLTRDLY | 0.04 |
| HVLTRT1 | 0.04 |
| NP2DRES | 0.04 |
| NP3RTCON | 0.04 |
| NP2SALV | 0.04 |
| DRMFIGHT | 0.04 |
| DRMAGRAC | 0.04 |
| VLTVEG | 0.04 |
| NP3PRSPR | 0.04 |
| NP3POSTR | 0.03 |
| NP3RIGRU | 0.03 |
| HVLTRT2 | 0.03 |
| NP2RISE | 0.03 |
| a_state | 0.03 |
| a_trait | 0.03 |
| VLTFRUIT | 0.03 |
| NP3FTAPL | 0.03 |
| ESS7 | 0.03 |
| NP3RIGLU | 0.03 |
| HVLTREC | 0.02 |
| NP2SPCH | 0.02 |
| LNS_TOTRAW | 0.02 |
| HVLTRT3 | 0.02 |
| NP3PRSPL | 0.02 |
| NP3TTAPL | 0.02 |
| NP2HWRT | 0.02 |
| DRMVERBL | 0.02 |
| MCATOT | 0.02 |
| gastrointestinal_up | 0.02 |
| CN2RSP | 0.02 |
| total | 0.02 |
| MCAVFNUM | 0.02 |
| JLO_TOTRAW | 0.01 |
| NP3HMOVR | 0.01 |
| NP1URIN | 0.01 |
| NP1CNST | 0.01 |
| NP3LGAGL | 0.01 |
| PN3RIGRL | 0.01 |

| clinical feature importance (Model: baseline+year1) → | baseline | year1 |
|---|---|---|
| NHY | 0.89 | 0.16 |
| NP3FACXP | 0.35 | 0.14 |
| NP3BRADY | 0.15 | 0.23 |
| NP2TRMR | 0.09 | 0.06 |
| urinary | 0.09 | 0.03 |
| NP3FTAPR | 0.09 | 0.02 |
| SDMTOTAL | 0.06 | 0.08 |
| NP3RIGRU | 0.06 | 0.04 |
| NP3RTCON | 0.05 | 0.12 |
| LNS_TOTRAW | 0.04 | 0.03 |
| NP3TTAPR | 0.04 | 0.02 |
| NP1SLPD | 0.04 | 0.02 |
| HVLTRT1 | 0.04 | 0.03 |
| gastrointestinal_down | 0.04 | 0.02 |
| VLTANIM | 0.04 | 0.03 |
| NP3GAIT | 0.04 | 0.08 |
| HVLTRT2 | 0.03 | 0.04 |
| NP2RISE | 0.03 | 0.01 |
| a_trait | 0.03 | 0.04 |
| NP3HMOVL | 0.03 | 0.05 |
| NP3HMOVR | 0.03 | 0.03 |
| VLTVEG | 0.03 | 0.04 |
| NP2HWRT | 0.03 | 0.06 |
| MCATOT | 0.03 | 0.10 |
| a_state | 0.03 | 0.03 |
| NP2DRES | 0.03 | 0.01 |
| MCAVFNUM | 0.03 | 0.03 |
| HVLTRDLY | 0.02 | 0.03 |
| VLTFRUIT | 0.02 | 0.02 |
| DRMAGRAC | 0.02 | 0.02 |
| NP3TTAPL | 0.02 | 0.02 |
| PN3RIGRL | 0.02 | 0.03 |
| NP2HOBB | 0.02 | 0.09 |
| HVLTRT3 | 0.02 | 0.03 |
| NP1LTHD | 0.02 | 0.00 |
| NP3RIGLU | 0.02 | 0.07 |
| NP1URIN | 0.02 | 0.01 |
| NP2SALV | 0.02 | 0.01 |
| HVLTREC | 0.02 | 0.01 |
| NP3POSTR | 0.02 | 0.02 |
| NP1SLPN | 0.02 | 0.01 |
| ESS5 | 0.02 | 0.06 |
| NP3RTALU | 0.01 | 0.01 |
| total | 0.01 | 0.04 |
| delayed_recall | 0.01 | 0.04 |
| gastrointestinal_up | 0.01 | 0.13 |
| DRMFIGHT | 0.01 | 0.01 |
| NP3FTAPL | 0.01 | 0.05 |
| NP2SPCH | 0.01 | 0.03 |
| NP3SPCH | 0.01 | 0.03 |

Table S3(b). Summary of clinical parameters with significant contributions to the prediction models. Table lists significantly contributing clinical parameters based on baseline examination tests (BL) or based on baseline with year-1 (BL + Y1) test items. Abbreviations: EPS, Epworth Sleepiness Scale; HVLT, Hopkins Verbal Learning Test; LNS, Letter-Number Sequencing; MDS- UPDRS, Movement Disorder Society Revision of the Unified Parkinson's Disease Rating Scale; MoCA, Montreal Cognitive Assessment; RBDSQ, REM Sleep Behavior Disorder Screening Questionnaire; QUIP, Questionnaire for Impulsive-Compulsive Disorders in Parkinson's Disease; SCOPA-AUT, Assessment of Autonomic Dysfunction; STAI, State-Trait Anxiety Inventory.

| **Clinical Scales** | **Model** | |
|---|---|---|
| **Specific Test Item(s) (*Parameter Name*)** | **BL** | **BL+Y1** |
| **MDS-UPDRS Part 1** | | |
| 1.7 Sleep problems (*NP1SLPN*) | + | + |
| 1.8 Daytime Sleepiness (*NP1SLPD*) | + | + |
| 1.10 Urinary Problems (*NP1URN*) | + | + |
| 1.11 Constipation problems (*NP1CNST*) | + | - |
| 1.12 Lightheadedness on Standing (*NP1LTHD*) | - | + |
| **MDS-UPDRS Part 2** | | |
| 2.1 Speech (*NP2SPCH*) | + | + |
| 2.2 Saliva and drooling (*NP2SALV*) | + | + |
| 2.5 Dressing (*NP2DRES*) | + | + |
| 2.7 Handwriting (*NP2HWRT*) | + | + |
| 2.8 Doing hobbies and other activities (*NP2HOBB*) | + | + |
| 2.10 Tremor (*NP2TRMR*) | + | + |
| 2.11 Getting out of bed, car, deep chair (*NP2RISE*) | + | + |
| **MDS-UPDRS Part 3** | | |
| 3.1 Speech (*NP3SPCH*) | - | + |
| 3.2 Facial expression (*NP3FACXP*) | + | + |
| 3.3a Rigidity – neck (*NP3RIGN*) | + | - |
| 3.3b Rigidity – RUE (*NP3RIGRU*) | + | + |
| 3.3c Rigidity – LUE (*NP3RIGLU*) | + | + |
| 3.3d Rigidity – RLE (*NP3RIGRL*) | + | + |
| 3.4a Finger tapping right hand (*NP3FTAPR*) | + | + |
| 3.4b Finger Tapping Left Hand (*NP3FTAPL*) | + | + |
| 3.5a Hand movements - Right Hand (*NP3HMOVR*) | + | + |
| 3.5b Hand movements – left hand (*NP3HMOVL*) | - | + |
| 3.6a Pronation-supination – right hand (*NP3PRSPR*) | + | - |
| 3.6b Pronation-supination – left hand (*NP3PRSPL*) | + | - |
| 3.7a Toe tapping - Right foot (*NP3TTAPR*) | - | + |
| 3.7b Toe tapping – left foot (*NP3TTAPL*) | + | + |
| 3.8b Leg agility – left leg (*NP3LGAGL)* | + | - |
| 3.10 Gait (*NP3GAIT*) | - | + |
| 3.13 Posture (*NP3POSTR*) | + | + |
| 3.14 Global Spontaneity of movement (*NP3BRADY*) | + | + |
| 3.17b Rest tremor amplitude- LUE (*NP3RTALU*) | - | + |
| 3.18 Constancy of rest (*NP3RTCON*) | + | + |
| 3.21 Hoehn and Yahr stage (*NHY*) | + | + |
| **MoCA** | | |
| Verbal fluency (*MCAVFNUM*) | + | + |
| MoCA total score (*MCATOT*) | + | + |

| | | | |
|---|---|---|---|
| **HVLT** | | | |
| | Immediate Recall Trial 1 (*HVLTRT1*) | + | + |
| | Immediate Recall Trial 2 (*HVLTRT2*) | + | + |
| | Immediate Recall Trial 3 (*HVLTRT3*) | + | + |
| | Delayed Recall (*HVLTRDLY*) | + | + |
| | Recognition (*HVLTREC*) | + | + |
| **LNS** | | | |
| | LNS-Sum questions 1-7 (*LNS_TOTRAW*) | + | + |
| **BENTON** | | | |
| | Judgment line of action total raw score (*JLO_TOTRAW*) | + | - |
| **RBDSQ** | | | |
| | Dreams frequently have aggressive or action-packed content (*DRMAGRAC*) | + | + |
| | In my dreams: speaking, shouting, swearing (*DRMVERBL*) | + | - |
| | In my dreams: sudden limb movements (*DRMFIGHT*) | + | + |
| **EPS** | | | |
| | Lying down to rest in the afternoon (*ESS5*) | - | + |
| | Sitting quietly after a lunch (*ESS7*) | + | - |
| **SCOPA-AUT** | | | |
| | Difficulty retaining urine + involuntary loss of urine + stream of urine been weak + pass urine at night + urine your bladder was not completely empty + urine again within 2 hours of the previous time (*urinary*) | + | + |
| | Had difficulty swallowing or have choked + Has saliva dribbled out of your mouth + Has food become stuck in your throat (*gastrointestinal_up*) | + | + |
| | Have feeling during meal that you were full very quickly + Had problems with constipation + Had to strain hard to press stools + Had involuntary loss of stools (*gastrointestinal_down*) | + | + |
| **Semantic Fluency** | | | |
| | Total number of animals (*VLTANIM*) | + | + |
| | Total number of vegetables (*VLTVEG*) | + | + |
| | Total number of fruits (*VLTFRUIT*) | + | + |
| **STAI** | | | |
| | Anxiety state score (*a_state*) | + | + |
| | Anxiety trait score (*a_trait*) | + | + |
| **SDM** | | | |
| | Total symbol digit modalities test (*SDMTOTAL*) | + | + |
| **Neuro Cranial** | | | |
| | Abnormality in Cranial Nerves (*CN2RSP*) | + | - |
| **Geriatric** | | | |
| | Geriatric depression total score (*total*) | + | + |

### Model Interpretation

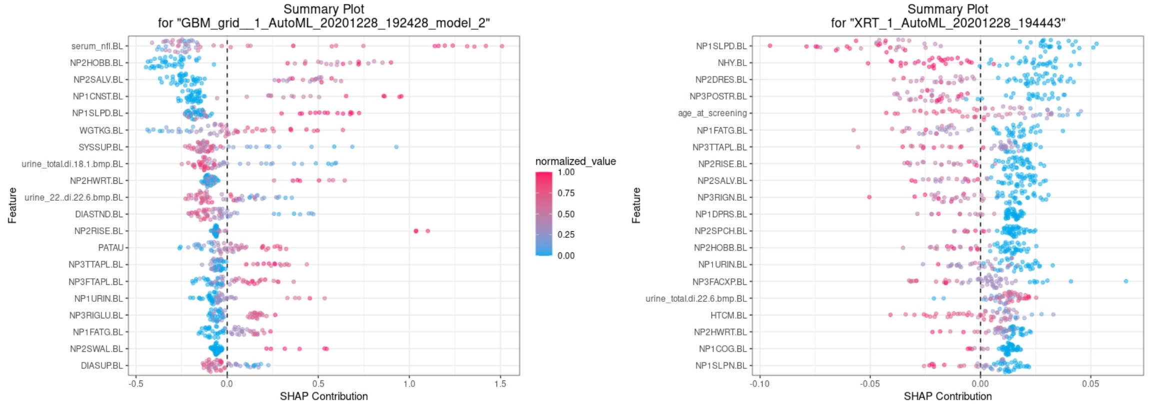

Fig S7: Clinical features influencing Parkinson's Disease progression class. Panels A and B from left to right. Detailed view of influence of top features for Higher PD progression class i.e. PDvec3 (Left) and lower PD progression class i.e. PDvec1 (Right). Higher value on the horizontal axis represents higher probability of a PD patient belonging to the PDvec3 class (left) and PDvec1 class (right).

SHAP is an unified approach to explain the output of any supervised machine learning model. It assigns an importance value to every feature based on Shapley values. In addition, it generates the impact of each feature on the model's output i.e. the class probability for classification algorithms. **Figure S7a** shows the impact score (SHAP contribution) on the probability of PDvec3 class. We see that a higher *serum_nfl* score corresponds to the increase in the probability of a patient belonging to the PDvec3 class. Similarly, higher scores on other symptomatic features related to hobby, sleep are among the top features that can differentiate between PD_h and other classes. **Figure S7b** shows the behavior of the model for PDvec1 class. The top features include sleeping behavior, Hoehn and Yahr stage score and the posture stability with probability of lower progressive class increases with increase in scores for these features. Younger PD patients at screening are expected to show lower PD progression as compared to the older patients. **Figure S8** shows the top 20 features involved in classifying PD progressive subtypes.

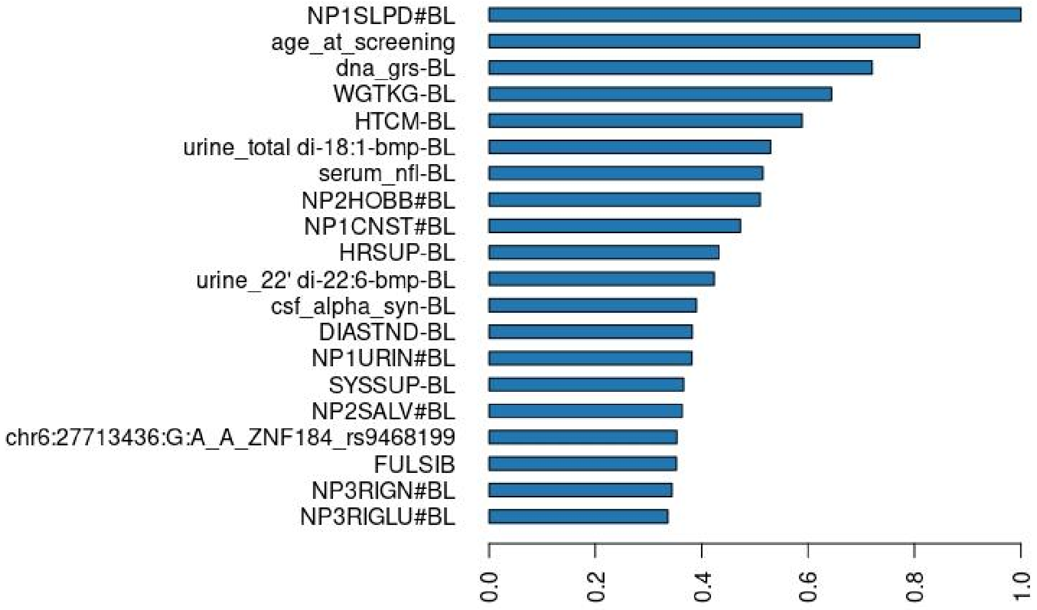

Figure S8: Shows the top-20 features with their importance score for the classification of different PD progression subtypes.

#### Change in diagnosis status

The clinical condition of patients can deteriorate, stay the same, or rarely gets better with time. As the study progresses and more information becomes available about the disease manifestation, the patient's diagnosis will be updated. We looked at cases where their clinical diagnosis were updated in the PPMI study. **Figure S9**, shows the trajectory of two patients initially diagnosed as PD in the progression space. We can observe that the patient whose status has changed from PD to dementia has much worse condition along the Cognitive dimension. The other patient whose status changed from PD to multiple system atrophy has shown more decline along the motor dimension.

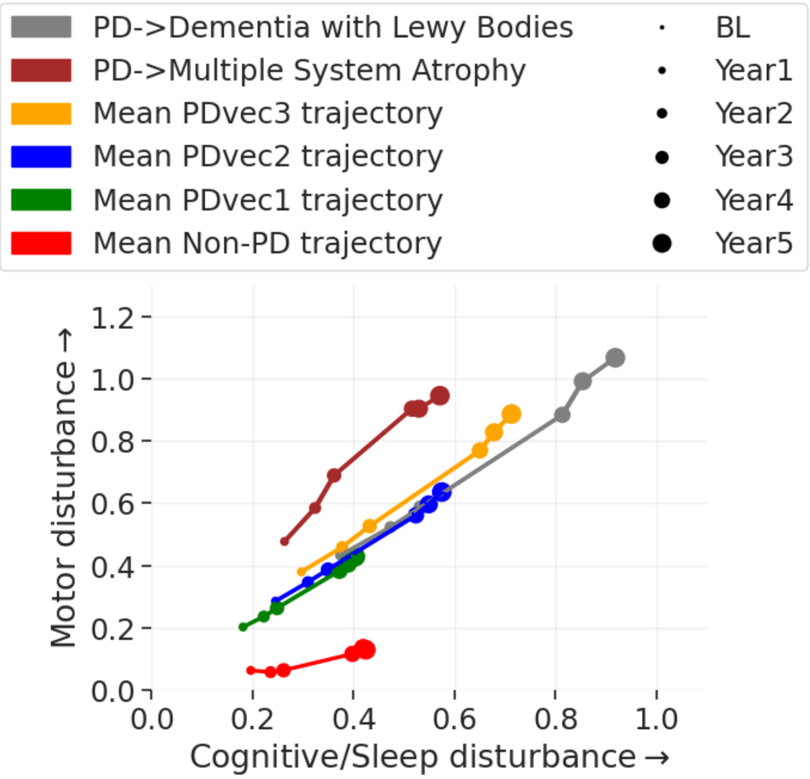

Fig S9. Shows the trajectory of two PD patients in the two dimensional progression space whose status has changed from their recruitment category in PPMI cohort. It also shows the average trajectory of PD subtypes and Non-PD subjects. The marker size corresponds to time from baseline.

**Table S4** below details baseline and follow-up differences in Nfl across the predicted progression vector classes. Here we show significant differences between slow and faster progressors not only in the measures of Nfl itself but in the slope of change.

Table S4. Shows the longitudinal changes in serum Nfl levels over 5 years for three subtypes. We used a statistical t-test between [a]PDvec1 vs. PDvec2 and [b]PDvec1 vs. PDvec3 to compare the means of slope and serum Nfl levels at different points in time.

| **Outcome** | **PDvec1** | **PDvec2** | **PDvec3** |
|---|---|---|---|
| *Δ serum Nfl level per year Mean [SD]* | 1.31 [2.36] | 1.48 [2.76] | 2.91 [4.23] |
| *t-test P-value [t-statistic]* | - | 0.6196[a] [-0.50] | 0.0025[b] [-3.07] |
| *Baseline Mean [SD]* | 11.54 [5.84] | 11.77 [5.21] | 15.42 [6.66] |
| *t-test P-value [t-statistic]* | - | 0.7576[a] [-0.31] | 0.0004[b] [-3.63] |

| | | | |
|---|---|---|---|
| *At end of year1 Mean [SD]* | 11.72 [5.05] | 13.69 [10.42] | 15.72 [6.38] |
| *t-test P-value [t-statistic]* | - | 0.086[a] [-1.73] | 0.0002[b] [-3.86] |
| *At end of year2 Mean [SD]* | 13.34 [8.09] | 13.85 [7.17] | 17.09 [7.91] |
| *t-test P-value [t-statistic]* | - | 0.6321[a] [-0.48] | 0.0138[b] [-2.49] |
| *At end of year3 Mean [SD]* | 14.16 [8.98] | 14.76 [8.98] | 19.87 [8.99] |
| *t-test P-value [t-statistic]* | - | 0.5761[a] [-0.56] | 0.0006[b] [-3.51] |
| *At end of year5 Mean [SD]* | 16.72 [13.51] | 18.02 [12.36] | 26.75 [20.41] |
| *t-test P-value [t-statistic]* | - | 0.4435[a] [-0.77] | 0.0003[b] [-3.73] |

### Supplementary references

None.
